# Flagellar dynamics reveal fluctuations and kinetic limit in the *Escherichia coli* chemotaxis network

**DOI:** 10.1101/2020.03.23.003962

**Authors:** Roshni Bano, Patrick Mears, Ido Golding, Yann R. Chemla

## Abstract

The *Escherichia coli* chemotaxis network, by which bacteria modulate their random run/tumble swimming pattern to navigate their environment, must cope with unavoidable number fluctuations (“noise”) in its molecular constituents like other signaling networks. The probability of clockwise (CW) flagellar rotation, or CW bias, is a measure of the chemotaxis network’s output, and its temporal fluctuations provide a proxy for network noise. Here we quantify fluctuations in the chemotaxis signaling network from the switching statistics of flagella, observed using time-resolved fluorescence microscopy of individual optically trapped *E. coli* cells. This approach allows noise to be quantified across the dynamic range of the network. Large CW bias fluctuations are revealed at steady state, which may play a critical role in driving flagellar switching and cell tumbling. When the network is stimulated chemically to higher activity, fluctuations dramatically decrease. A stochastic theoretical model, inspired by work on gene expression noise, points to CheY activation occurring in bursts, driving CW bias fluctuations. This model also shows that an intrinsic kinetic ceiling on network activity places an upper limit on activated CheY and CW bias, which when encountered suppresses network fluctuations. This limit may also prevent cells from tumbling unproductively in steep gradients.

## Introduction

A common feature of living organisms is their ability to sense environmental signals and respond to these signals by modifying their behavior. This ability is enabled by signal transduction, which converts environmental inputs into behavioral outputs^1^. Studies have shown that signaling networks must operate in the presence of noise not only in their inputs but also in the circuit components themselves^2,3^. Until recently, it was thought that signaling networks had evolved to be robust against undesirable noise^4–7^. More recent studies have shifted to understanding the role of noise in regulating and fine-tuning biological function^8,9^. For example, noise resulting from transcription/translation has been measured precisely and shown to play a critical role in networks for gene expression and regulation^10–12^. In contrast, studying noise resulting from signaling networks at the post-translation level has proven more challenging.

One extensively studied signaling system is the chemotaxis network of *E. coli*, which cells use to navigate their environment^13–17^. *E. coli* cells swim in a random walk consisting of “runs”— during which their flagella rotate counter-clockwise (CCW)—and “tumbles”—during which one or more flagella rotate clockwise (CW)^18–21^. Changing environmental conditions are sensed through a two-component signaling system comprising receptors that bind extracellular ligands and a kinase, CheA, that transfers its phosphoryl group onto downstream effectors^22^. The response regulator, CheY, when phosphorylated by the receptor kinase complex to CheY-P, binds to the flagellar motor and increases the probability of CW motor rotation^23–25^. The phosphatase CheZ carries out the opposite reaction, maintaining a dynamic equilibrium between CheY and CheY-P (**Figure 1a**)^26,27^. Additionally, the methyltransferase CheR and the methylesterase CheB covalently modify receptors, and the resulting modulation of CheA activity affects CheY-P levels over the timescales of chemotactic adaptation^28–30^. While the population-averaged relationships between these signaling protein concentrations and cellular motility in *E. coli* are now well understood^13–16,24^, the role of fluctuations in modulating cell behavior remains comparatively unexplored.

**Figure 1.**
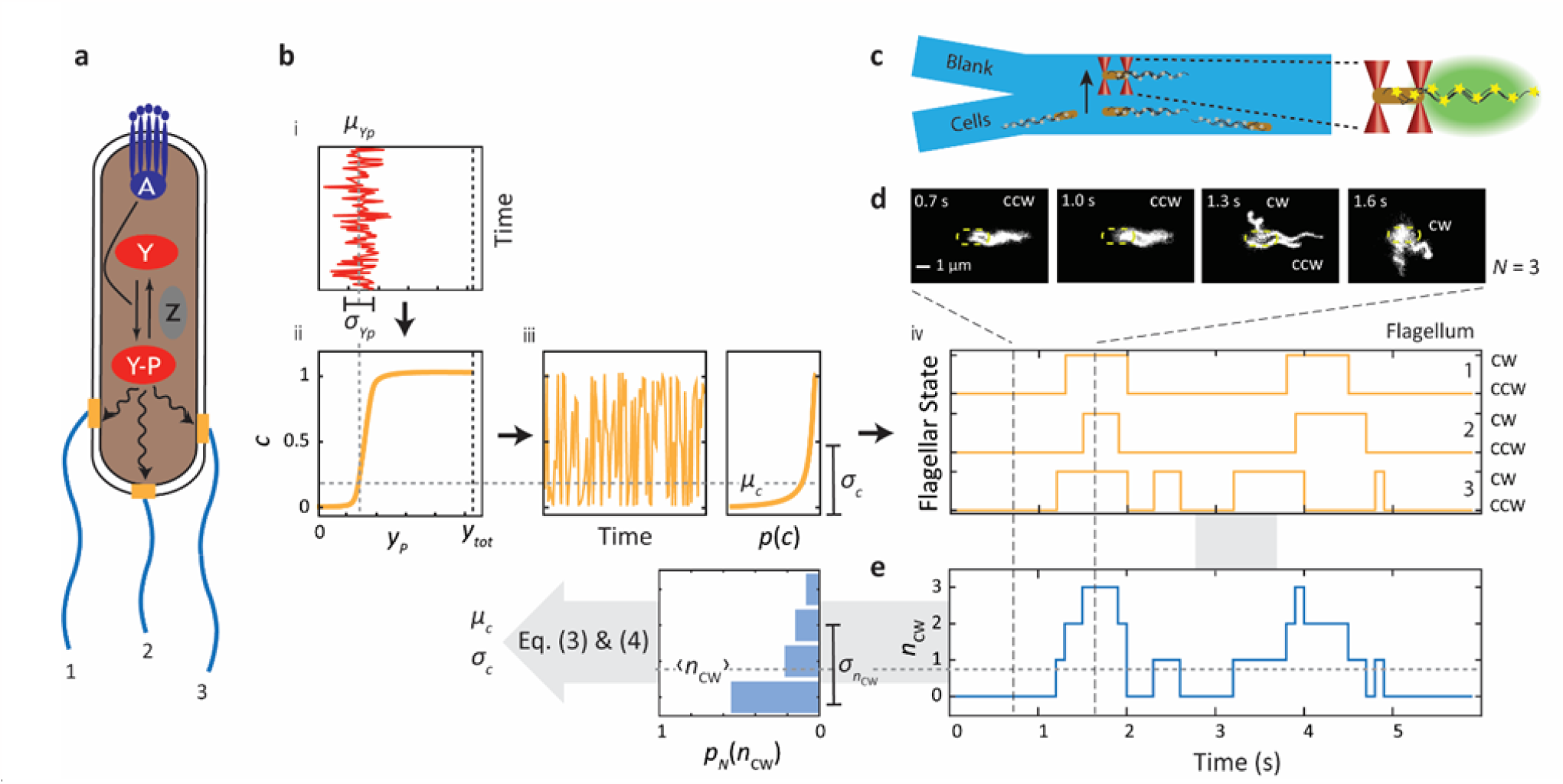
Estimating the temporal motor bias fluctuations from flagellar dynamics of single *E. coli* cells. **a**. Schematic of a multi-flagellated *E. coli* cell showing key reactions of the chemotaxis signaling network. CheY is phosphorylated to CheY-P by the receptor kinase complex (CheA; dark blue); CheY-P is dephosphorylated by the phosphatase CheZ (gray). CheY-P binding to flagellar motors (mustard) causes a switch from CCW to CW rotation, which leads to cell tumbling. **b**. Temporal fluctuations in [CheY-P] due to chemotaxis network dynamics and their effect on flagellar switching statistics. (i) [CheY-P] temporal fluctuations. (ii) Switch-like response of motor CW bias, *c*, to [CheY-P]. (iii) Resulting temporal fluctuations in motor bias *c*(*t*) and its distribution *p*(*c*), characterized by the mean *μ*_*c*_ and standard deviation *σ*_*c*_. (iv) CCW/CW rotation state of three independent flagellar motors corresponding to the instantaneous motor bias *c*. **c**. Schematic of two-channel laminar flow chamber for optical trap assay. A single cell with fluorescently labelled flagella (gray stars) is captured from the bottom channel (“Cells”), aligned along the flow between two optical traps (red cones), moved to the upper channel (“Blank”), and its fluorescent flagella (yellow stars) imaged by stroboscopic slim-field microscopy (green). **d**. Fluorescence data trace from representative trapped cell with *N* = 3 flagella. Top, still images of fluorescent flagella, with position of the unlabeled cell body approximately indicated by the dashed yellow line. CW and CCW rotating flagella are labelled where clearly visible. Middle, CCW/CW rotation state of each flagellum vs. time (mustard). Bottom, corresponding number *n*_CW_ of CW flagella vs. time (light blue). **e**. Probability distribution *p*_*N*_(*n*_CW_), determined experimentally from d (light blue bars). The mean *μ*_*c*_ and standard deviation *σ*_*c*_ of the motor bias are estimated from the parameters of the distribution *p*_*N*_(*n*_CW_) using Eqs. (3) and (4).

Noise in the chemotaxis network was first inferred by Korobkova et al.^31^ from long-term measurements of individual rotating flagellar motors. Inside the cell, the CheY-P concentration [CheY-P] fluctuates in time due to noise sources upstream in the signaling network (**Figure 1b, i**). These fluctuations in turn affect downstream behavior. The motor CW bias, i.e. the probability of CW rotation of a single flagellar motor, has an ultrasensitive switch-like dependence^25^ on the intracellular [CheY-P] (**Figure 1b, ii**), resulting in fluctuations in motor rotational state (**Figure 1b, iii**) and in the cell’s swimming behavior. Noise in the *E. coli* chemotaxis signaling network is thought to have functional consequences for sensing and navigation in complex natural microenvironments. It has been proposed to allow cells to sample 3-D space more effectively^31,32^, enhance chemotactic drift up a gradient^33–35^, and synchronize flagellar switching to mitigate differences in swimming behavior among cells with different numbers of flagella^36,37^.

Here, we use optical trapping coupled with time-resolved fluorescence microscopy^37,38^ to infer temporal network fluctuations from the statistics of flagellar switching in individual multi-flagellated *E. coli* cells. Analyzing cells at steady state, we determine the mean and fluctuations in CW bias, which reveal that the network is poised at a low CW bias, favoring CCW flagellar rotation/running, and that transient fluctuations drive CW flagellar rotation/tumbling. Previous studies using flagellar motor behavior^31,39^ or fluorescence-based reporters^40,41^ to infer noise in the chemotaxis network have been limited to long timescales (>10 s) due to temporal averaging. However, studies of flagellar tracking show that short timescale fluctuations are important to explain observations of correlated flagellar switching^36,42^. In contrast to most earlier approaches, our method measures fluctuations on shorter timescales (∼3 s), comparable to individual run and tumble durations. We exploit this feature to track carefully the time evolution of signaling noise in cells responding to a stimulus that increases CheA activity. This, in turn, allows us to measure network fluctuations across the accessible range of motor CW biases. We find that fluctuations are significantly reduced when the motor CW bias is increased away from its steady-state value. A simple stochastic model in which noise in signaling arises from burst-like fluctuations in CheA activity recapitulates all of our experimental results and suggests that the decrease in fluctuations upon stimulation is due to a kinetic ceiling on the CheA activity.

## Results

### Flagellar switching statistics reveal temporal network fluctuations

We inferred the temporal fluctuations in the chemotaxis network from the dynamics of flagellar switching in individual cells. We imaged fluorescently labeled flagella on individual, swimming cells to determine their CCW/CW rotation state, using an optical trapping assay described previously^37,38,43^ (see **Materials and Methods**). Briefly, *E. coli* cells were injected into a laminar flow chamber containing two parallel channels. A single cell was captured using a dual beam optical trap from one channel containing many cells (labeled “cells” in **Figure 1c**) and moved for observation to a second channel containing no other cells (labeled “blank” in **Figure 1c**). The light scattered by the optically trapped cell was used to monitor the run/tumble swimming behavior as described previously (data not shown, see Ref. [38]). Simultaneously, high-speed, epi-fluorescent stroboscopic imaging^37^ was used to monitor the rotation state of all the individual flagella on the cell at high temporal resolution (0.1 s) and long-time duration (ranging from ∼15 to 40 s, until the flagella photobleached) (**Figure 1d**, top). Following Darnton et al.^21^ and our previous work^39^, the CCW/CW rotation state of each flagellum was visually identified by its different helical waveforms (see **Materials and Methods**).

We next inferred network fluctuations making use of the statistics of switching in multi-flagellated cells. For a cell with *N* flagellar motors, the probability that *n*_*cw*_ rotate in a CW direction, *p*_*N*_ (*n*_*cw*_), is equal to (see **Materials and Methods**):

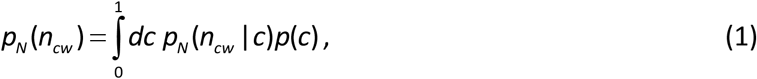

where *c* is the instantaneous motor CW bias, or probability that a motor rotates CW, *p*_*N*_ (*ncw* |*c*) is the conditional probability that *n*_*cw*_ out of *N* motors are in the CW state given a certain *c*, and *p*(*c*) is the distribution of CW biases (**Figure 1b, iii**). Previous work in which flagellar motor switching was decoupled from the chemotaxis signaling network shows that the motors independently sense the intracellular CheY-P concentration^37^. Thus, *p*_*N*_ (*n*_*cw*_ |*c*), represents the conditional probability of the collective CW state of the flagella motors and is given by the binomial distribution:

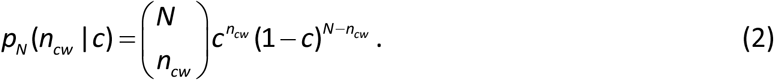

Since the conditional probability is known, Eq. (1) in principle allows one to connect the probability *p*_*N*_ (*n*_*cw*_) —which can be determined from measurements of the CCW/CW rotational state of each flagellum of a cell with *N* flagellar motors—to the distribution of CW biases *p*(*c*). In practice, a solution to Eq. (1) is not unique, but an increasing number of moments of *p*(*c*) can be determined as the number of flagella *N* increases. Specifically, from Eqs. (1) and (2), the mean and variance in the number of CW flagella *n*_*cw*_ can be shown to be (see **Materials and Methods**)

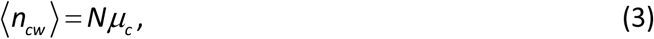

and

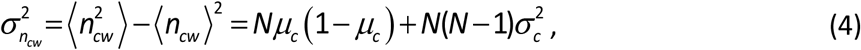

where *μ*_*c*_ and *σ*_*c*_ are the mean and standard deviation in CW bias. In Eq. (4) the first term represents the fluctuations in *p*_*N*_ (*n*_*cw*_) intrinsic to switching of *N* independent flagellar motors; the second term represents the additional contribution from fluctuations in CW bias. These expressions show that by analyzing cells with *N* ≥ 2 flagella, one can determine the mean and fluctuations in CW bias over a given time window.

Figure 1. shows the workflow to determine network fluctuations experimentally. Fluorescence tracking of the CCW/CW state of the individual flagella (**Figure 1d**, middle) is represented as a time trace of the number of CW flagella (**Figure 1d**, bottom). For a cell with *N* flagella, a histogram of this time trace was used to construct the probability distribution *p*_*N*_ (*n*_*cw*_) that *n*_*cw*_ out of *N* flagella rotate CW (**Figure 1e**), from which the mean and standard deviation in *n*_*cw*_ were calculated. Using Eqs. (3) and (4) the mean, *μ*_*c*_, and standard deviation, *σ*_*c*_, in CW bias for the cell were determined (**Figure 1e**).

### Cells at steady state exhibit large CW bias fluctuations

We first applied the above approach to free-swimming (i.e. unstimulated) cells for which the chemotaxis network is in a steady state. Fluorescence tracking data of CCW/CW flagellar states were obtained for cells of a strain wild-type for chemotaxis (HCB1660, henceforth referred to as “wild-type”; see **Materials and Methods**) with *N* = 2 to 5 flagella. Data traces tracking *n*_*cw*_ were pooled together from multiple cells with the same total number of flagella *N* to generate *p*_*N*_ (*n*_*cw*_) (**Figure 2a**, colored histograms; number of cells = 13, 15, 14, 9 for *N* = 2, 3, 4, 5 flagella, respectively). As we show below, pooling over the population of cells did not affect our results appreciably.

**Figure 2.**
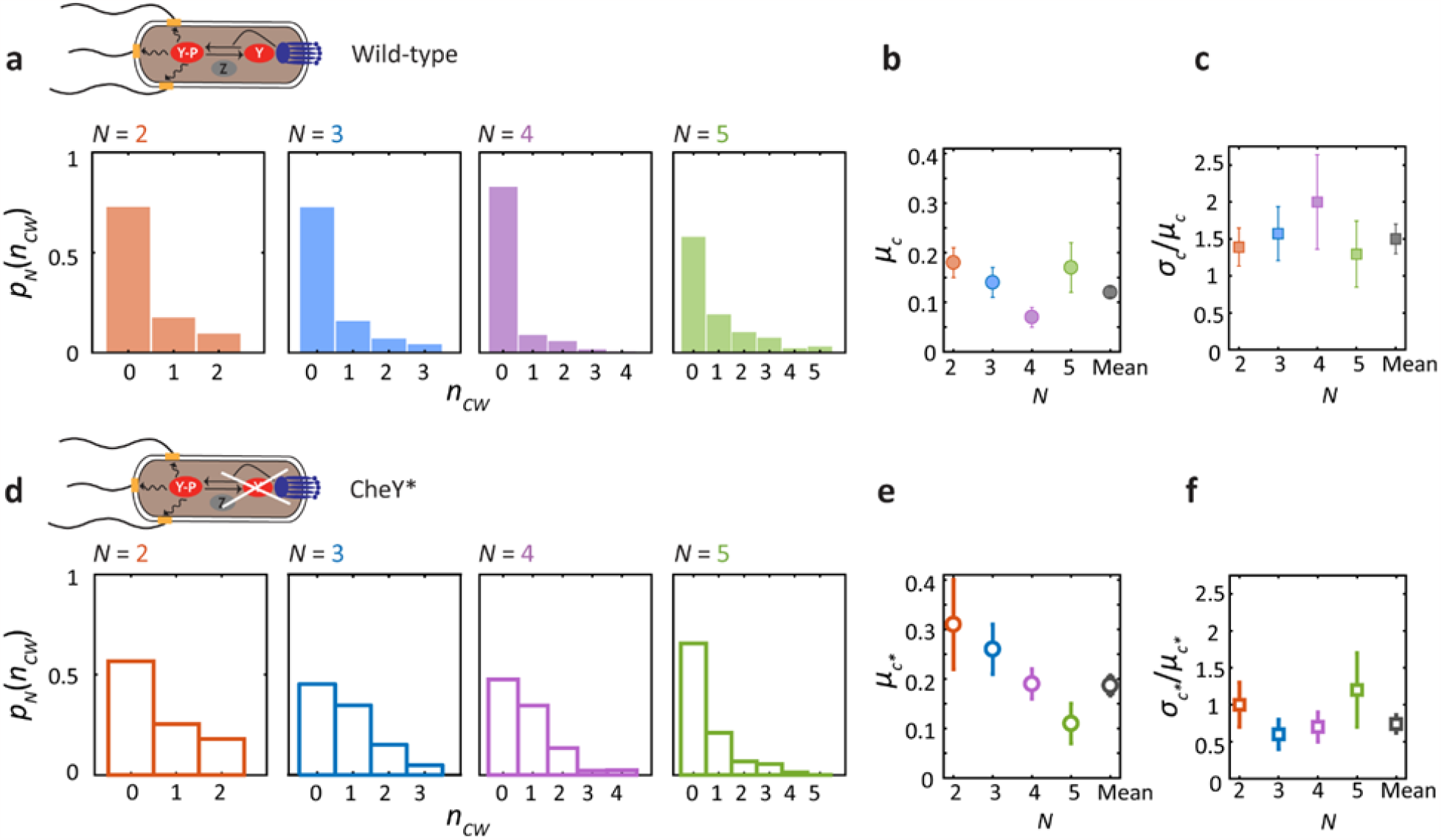
Motor bias fluctuations in the chemotaxis network at steady state. Measurement of flagellar dynamics and estimation of motor bias fluctuations in wild-type *E. coli* strain HCB1660 (a-c) and in “CheY*” mutant strain PM87 expressing constitutively active CheY^D13K^ (d-f). **a**. Experimentally determined probability distribution *p*_*N*_(*n*_CW_) that *n*_CW_ out of *N* flagella rotate CW, grouped by cells with the same total number of flagella *N* = 2, 3, 4, 5 (number of cells = 13, 15, 14, 9 respectively). **b**. Mean motor bias, *μ*_*c*_, determined separately for each *N* (colored filled circles) and the mean across *N* (gray filled circle). **c**. Coefficient of variation in motor bias, *σ*_*c*_/*μ*_*c*_, determined separately for each *N* (colored filled squares) and the mean across *N* (gray filled squares) **d-f**. Same as a-c for CheY* strain, showing experimental distributions (open bars) and estimated motor bias parameters (open circles and squares), grouped by cells with the same total number of flagella *N* = 2, 3, 4, 5 (number of cells = 13, 13, 9, 9 respectively).

**Figures 2b-c** shows the mean, *μ*_*c*_, and coefficient of variation (CV), *σ*_*c*_/*μ*_*c*_ in CW bias for each set of cells with different number of flagella *N*, as determined from Eqs. (3) and (4). Errors in *μ*_*c*_ and *σ*_*c*_/*μ*_*c*_ were estimated by bootstrapping over the population of cells with the same *N* (see **Materials and Methods**). The parameter values did not appear to depend strongly on *N*, consistent with our expectation that the number of flagella should not affect network fluctuations measurably. Thus, we also analyzed the data globally, carrying out a weighted average over all cells (number of cells = 51) to determine <*μ*_*c*_> = 0.12 ± 0.01 and <*σ*_*c*_/*μ*_*c*_> = 1.5 ± 0.2 (see **Materials and Methods**). A CV > 1 shows that the chemotaxis network in free-swimming wild-type cells exhibits a high degree of fluctuations.

We carried out several tests to confirm that the observed fluctuations in CW bias resulted from noise in signal transduction. First, we applied our method of analysis to a strain that expresses the constitutively active mutant protein CheY^D13K^, a mimic for CheY-P^24^ (strain PM87, henceforth referred to as “CheY*”; see **Materials and Methods**). Since CheY in this strain is decoupled from the upstream network components and the concentration of CheY^D13K^ is determined from gene expression levels, which should not fluctuate on the timescales of our experiments in nutrient-limited conditions^44^, we expected to observe smaller fluctuations in CW bias as compared to the wild-type strain. Repeating the above analysis, **Figure 2d** show the experimental distributions *p*_*N*_ (*n*_*cw*_). Values for the mean, standard deviation, and CV in CW bias are found in **Tables 3** and **4**. Comparing **Figs. 2b-c** and **2e-f**, the mean CW bias is similar for both strains (0.12 ± 0.01 and 0.19 ± 0.02, respectively), and, as expected, fluctuations in the CheY* strain (number of cells = 13, 13, 9, 9 for *N* = 2, 3, 4, 5 flagella, respectively) are reduced compared to the wild-type strain, yielding <*σ*_*c*_/*μ*_*c*_> = 0.7 ± 0.2 (total number of cells = 44).

**Table 1.**
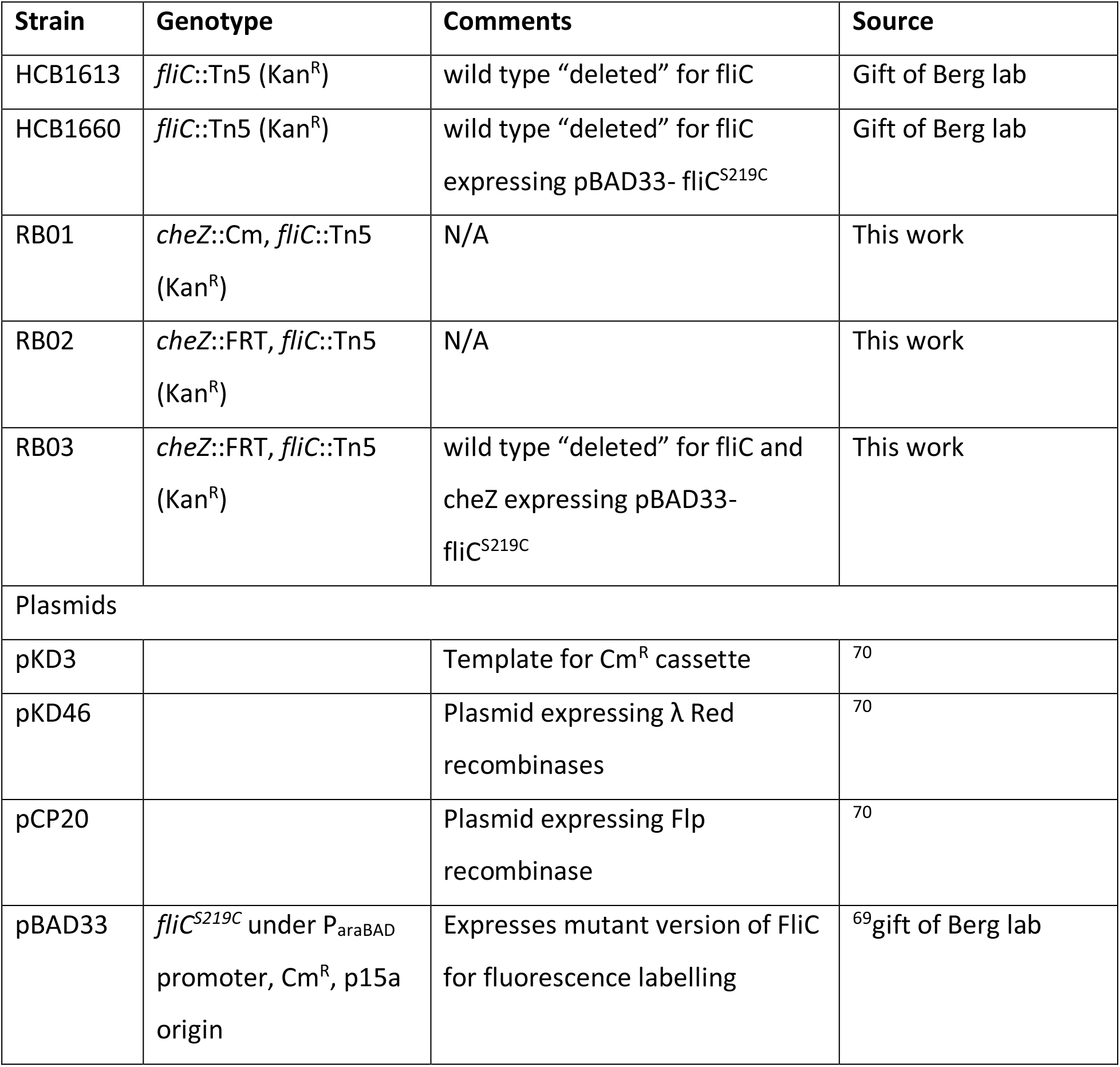
Strains and Plasmids used in this study.

**Table 2.**
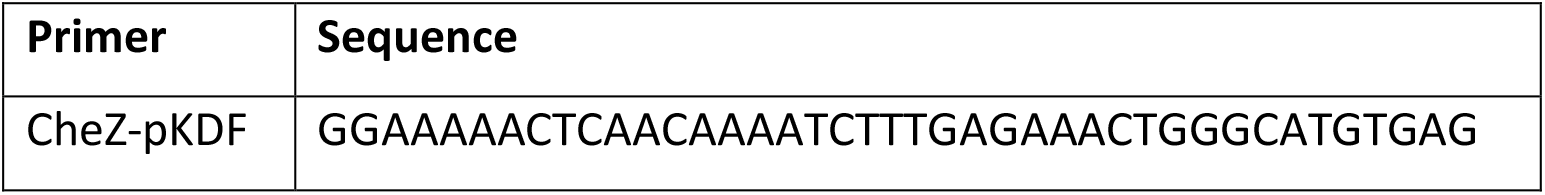
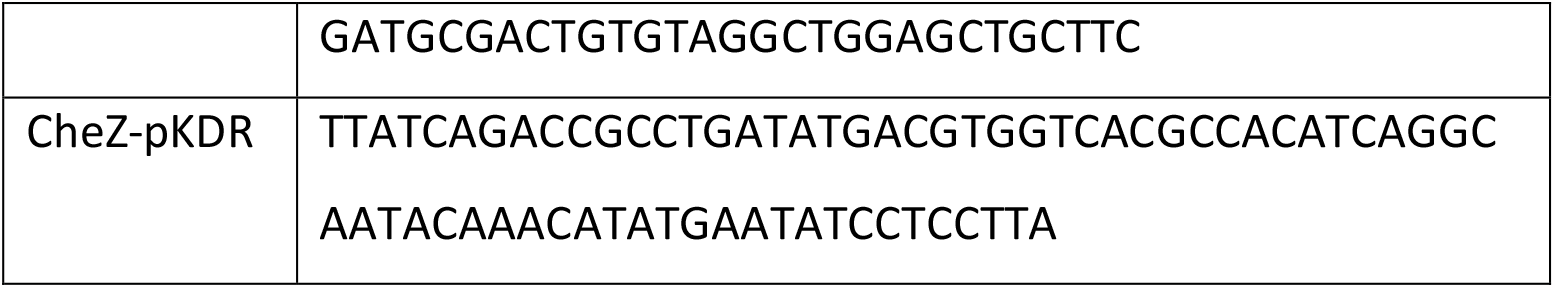
Primers used in this work.

**Table 3.** CW bias parameters for the wild type strain.

| $N$ | Mean CW bias,<br>$\mu_c$ | Standard deviation<br>in CW bias, $\sigma_c$ | Coefficient of Variation,<br>$\sigma_c/\mu_c$ |
| --- | --- | --- | --- |
| 2 | $0.18 \pm 0.03$ | $0.25 \pm 0.02$ | $1.4 \pm 0.3$ |
| 3 | $0.14 \pm 0.03$ | $0.22 \pm 0.02$ | $1.6 \pm 0.4$ |
| 4 | $0.07 \pm 0.02$ | $0.14 \pm 0.02$ | $2.0 \pm 0.6$ |
| 5 | $0.17 \pm 0.05$ | $0.22 \pm 0.03$ | $1.3 \pm 0.4$ |
| Global | $0.12 \pm 0.01$ | $0.20 \pm 0.01$ | $1.5 \pm 0.2$ |

**Table 4.** CW bias parameters for the mutant CheY* strain.

| $N$ | Mean CW bias,<br>$\mu_c$ | Standard deviation<br>in CW bias, $\sigma_c$ | Coefficient of Variation,<br>$\sigma_c/\mu_c$ |
| --- | --- | --- | --- |
| 2 | $0.31 \pm 0.09$ | $0.29 \pm 0.06$ | $1.0 \pm 0.3$ |
| 3 | $0.26 \pm 0.05$ | $0.17 \pm 0.03$ | $0.6 \pm 0.2$ |
| 4 | $0.19 \pm 0.03$ | $0.14 \pm 0.03$ | $0.7 \pm 0.2$ |
| 5 | $0.11 \pm 0.04$ | $0.13 \pm 0.03$ | $1.2 \pm 0.5$ |
| Global | $0.19 \pm 0.02$ | $0.16 \pm 0.02$ | $0.7 \pm 0.1$ |

Second, we tested whether the variance in CW bias could be due to cell-to-cell differences in mean CW bias rather than fluctuations arising in signal transduction. To test the contribution of cell-to-cell variation, we carried out the above analysis over individual cell traces and compared the wild-type and CheY* strains. As shown in **Figure 3a** and **3b**, while the mean CW bias, *μ*_*c*_, from individual cell distributions was similar for both strains, the CV, *σ*_*c*_/*μ*_*c*_, was significantly smaller for the CheY* strain as compared to the wild-type strain (*p* ≤ 0.05; determined using a Wilcoxon rank sum test), as expected. Errors in *μ*_*c*_ and *σ*_*c*_/*μ*_*c*_ for individual cells were again determined by bootstrapping (see **Materials and Methods**). Importantly, the coefficient of variation for individual wild-type cells averaged over all cells with different *N*, <*σ*_*c*_/*μ*_*c*_> ≈ 1.5, was close to that determined over the population. This indicates that cell-to-cell variation over the wild-type cell population does not contribute greatly to the variance in CW bias. In contrast, the CV was significantly smaller for individual cells of the CheY* strain; <*σ*_*c*_/*μ*_*c*_> ≈ 0.4, as compared to 0.7 over the population. We suspect cell-to-cell CheY^D13K^ expression differences may contribute to the estimated fluctuations in **Figures 2e-f**. Furthermore, we carried out a variance decomposition analysis to identify the contribution of cell-to-cell differences to the variance in CW bias measured over the cell population (see **Materials and Methods**). Consistent with the results above, we found that only 20% of the variance in CW bias for wild-type cells resulted from cell-to-cell differences, compared to 70% for CheY* cells. Our results thus show that we can attribute the observed variance in CW bias for wild-type cells to fluctuations due to signaling activity in individual cells.

**Figure 3.**
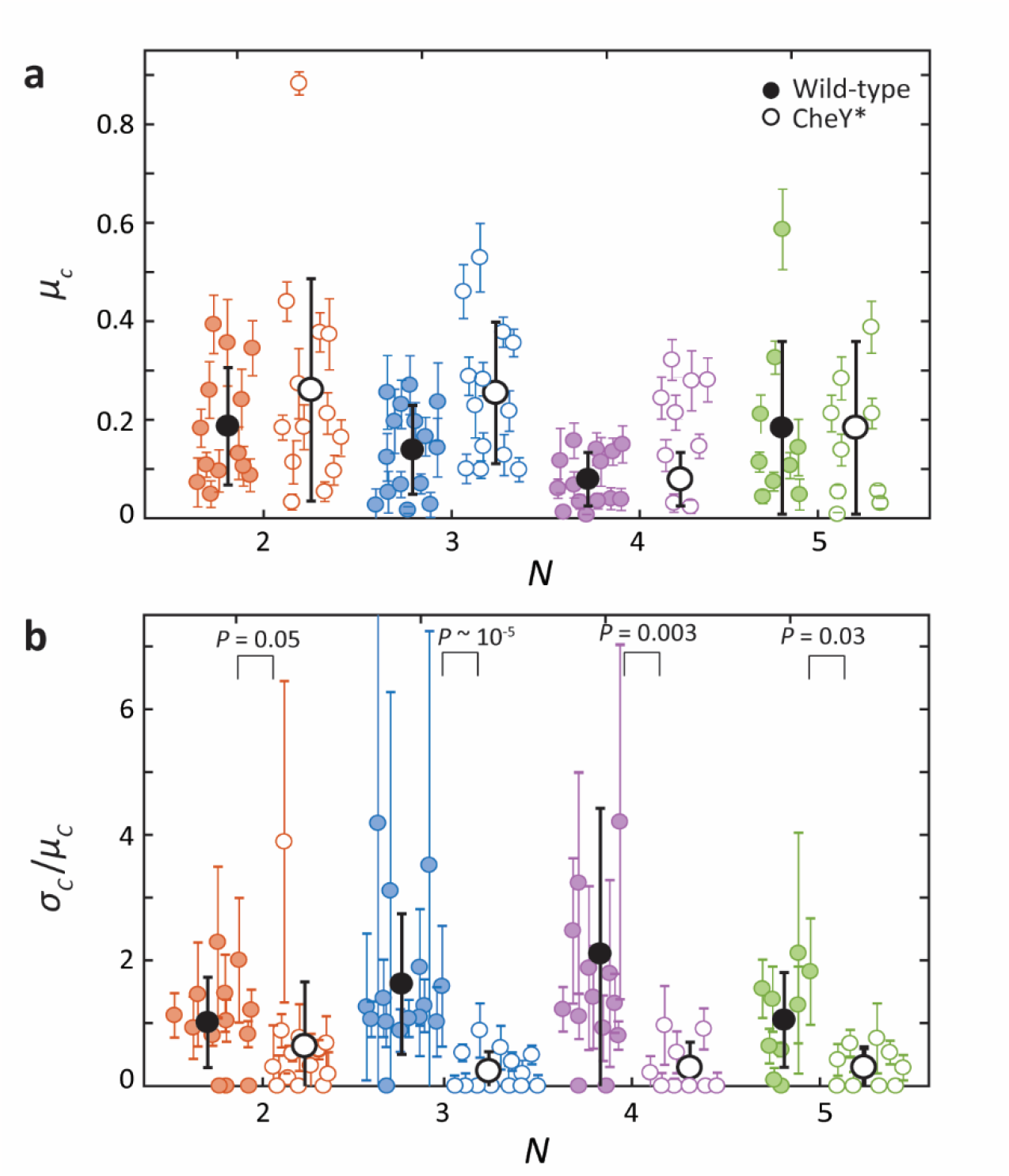
Estimate of the mean and coefficient of variation of motor bias in individual cells. **A**. Comparison of mean motor bias, *μ*_*c*_, determined for individual wild-type (colored circles) and CheY* cells (open circles), for different number of flagella *N*. **b**. Comparison of the coefficient of variation in motor bias, *σ*_*c*_/*μ*_*c*_, determined for individual wild-type (colored circles) and CheY* cells (open circles), for different *N*. Averages over each cell population are overlaid (black solid circles for wild-type, black open circles for CheY*); error bars represent standard deviation. P-values were determined using a Wilcoxon rank sum test.

Our method for determining network fluctuations allows us to make an important observation: the mean CW bias in free swimming cells is low (0.12), while the fluctuations in CW bias are very large (CV of 1.5). Thus the picture emerges of a network poised at low CW bias, corresponding to CCW flagellar rotation and running, with large fluctuations driving CW flagellar rotation and tumbling. This picture is consistent with simulations from Dufour et al.^45^ as well as experiments from Wong-Ng et al.^46^ showing that cells’ migration speed along chemical gradients is maximized at very low CW bias^47^. All these observations suggest that the network operates at a point to optimize chemotactic drift, and that fluctuations function to expand the allowed range of run/tumble behavior of the cell.

### Fluctuations decrease upon stimulating the network to increase CW bias

Having characterized fluctuations in the chemotactic network in the steady state, we next investigated them in a network stimulated by a perturbation. In response to a stepwise change in chemical environment, the intracellular [CheY-P] changes rapidly, and then reverts to its steady-state value through adaptation^48–50^. Applying a large stepwise decrease in attractant leads to a transient increase in [CheY-P] and in motor CW bias, followed by a decrease to the steady state^43,51^, allowing us to measure network fluctuations across the accessible range of motor CW biases.

We measured the response of individual cells to a decrease in concentration of α-methyl-DL-aspartate (MeAsp), a non-metabolizable analog of L-aspartate^52^ that acts as a chemoattractant^52,53^. We used a three-channel flow chamber (**Figure 4a**) to observe the effect of a step-down from 1 to 0 mM MeAsp on individual optically trapped wild-type cells. The step-down was applied by quickly moving a trapped cell from the middle channel of the flow chamber containing 1 mM MeAsp (**Figure 4a**; labeled “blank + attractant”) to the top channel containing no MeAsp (labeled “blank”; see **Materials and Methods**). The run-tumble behavior of the cell was monitored pre- and post-stimulus via the optical traps (see **Materials and Methods**), whereas the flagellar rotation state was measured by fluorescence imaging only post-stimulus.

**Figure 4.**
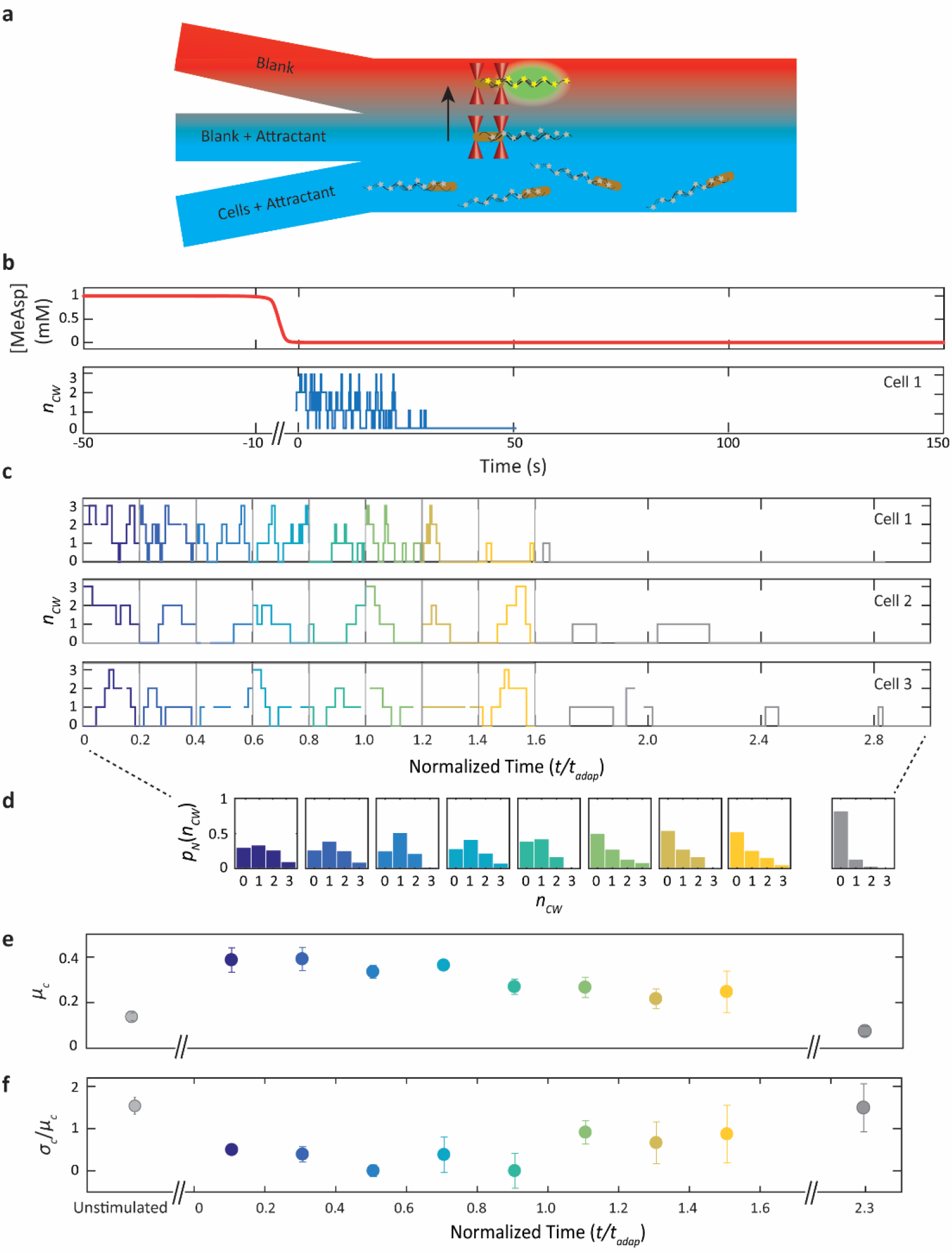
Motor bias fluctuations in the chemotaxis network upon stimulation. **a**. Schematic of three-channel laminar flow chamber for flagellar imaging of a cell experiencing a step down in attractant concentration. A single cell with fluorescently labelled flagella (gray stars) is captured from the bottom channel containing 1 mM MeAsp (“Cells + Attractant”) and aligned along the flow between two optical traps (red cones) in the middle channel (“Blank + attractant”). The trapped cell is then moved to the upper channel lacking MeAsp (“Blank”) and its fluorescent flagella (yellow stars) are imaged (green). **b**. Representative trace showing the response of “Cell 1” to a step-down stimulus. Top, schematic of MeAsp concentration over time, showing the step down from 1 to 0 mM. Data are not collected during the move from the middle to the top channel. Bottom, number *n*_CW_ of CW flagella over time after the move has finished at *t* = 0. **c-e**. Estimation of motor bias fluctuations in cells responding to a step-down stimulus. **c**. Traces from three representative “Cells 1, 2, 3” showing the number *n*_CW_ of CW flagella vs. time (normalized by each cell’s adaptation time *t*_*adap*_). Each trace is subdivided into windows of duration 0.2*t*_*adap*_, and data from each window is pooled across cells (for *N* = 3 flagella shown here, number of cells = 9). **d**. Distributions of the number of CW flagella *p*_*N*_(*n*_CW_) for each window (colored bars) **e**. Mean motor bias *μ*_*c*_ corresponding to each window and for unstimulated cells (gray) as comparison. **f**. Coefficient of variation in motor bias *σ*_*c*_/*μ*_*c*_ corresponding to each window, and for unstimulated cells.

**Figure 4b** shows a representative measurement of a cell with 3 flagella undergoing a step-down stimulus, where we define *t* = 0 as the time at which the trapped cell has finished moving to the channel with no MeAsp. In response, the cell exhibits the expected increase in tumbling immediately post-stimulus, followed by relaxation to the pre-stimulus steady state as the cell adapts (**Supplementary Figure S1**). As the probability of tumbling is directly related to the probability of CW flagellar rotation^37^, *n*_*cw*_ measured by fluorescence imaging post-stimulus also relaxes to lower values as the cell adapts (**Figure 4b**, blue trace). Since the motor bias is time-dependent in cells responding to a step-down, the time traces of *n*_*cw*_ for each cell were divided into time windows corresponding to 20% of the cell’s adaptation time, or ∼2.6 s on average (**Figure 4c**, colors corresponding to time window), over which the average tumble bias was constant to within ∼10-20% (see **Materials and Methods** for more details on windowing). All data from cells with the same total number of flagella (in **Figure 4c**, three representative cells with *N* = 3 flagella) were pooled in similarly constructed time windows and used to create histograms *p*_*N*_ (*n*_*cw*_) at each time (**Figure 4d**). The mean and CV in the CW bias were then estimated separately for each window (**Figure 4e-f**, colored data points). Relative to unstimulated cells (**Figure 4e**, gray data points for unstimulated wild-type cells with *N* = 3 flagella), the mean CW bias shifted to a higher value immediately after the stimulus, as expected for a step down. Interestingly, the CV in CW bias decreased (**Figure 4f**). We verified that the reduction in noise was due to a change in chemotaxis network dynamics after stimulation rather than due to windowing of the data; the standard deviations in CW bias in adapting cells were significantly different compared to those in steady-state cells over similarly sized 3-s time windows (*p* ≤ 0.05).

The time-dependent, post-stimulus mean CW bias *μ*_*c*_ and coefficient of variation *σ*_*c*_/*μ*_*c*_ for cells with *N* = 2, 3, 4 flagella are shown in **Figure 5a** and **5b** (colored filled circles; number of cells = 8, 9, 9), respectively. The corresponding steady-state mean and coefficient of variation obtained from **Figure 2** are also plotted for reference (for *t* < 0, colored open circles). As expected, the trends are similar for cells with different numbers of flagella. Thus, we also show these parameters determined globally for each time window by a weighted average across cells with different *N* (**Figure 5a** and **5b**; gray filled circles; see **Materials and Methods**). Consistent with **Figure 4e-f**, the mean CW bias increases immediately in response to the step-down stimulus and relaxes toward its steady-state, pre-stimulus value as the cells adapt, whereas the CV in CW bias falls post-stimulus, then gradually increases toward the pre-stimulus value.

**Figure 5.**
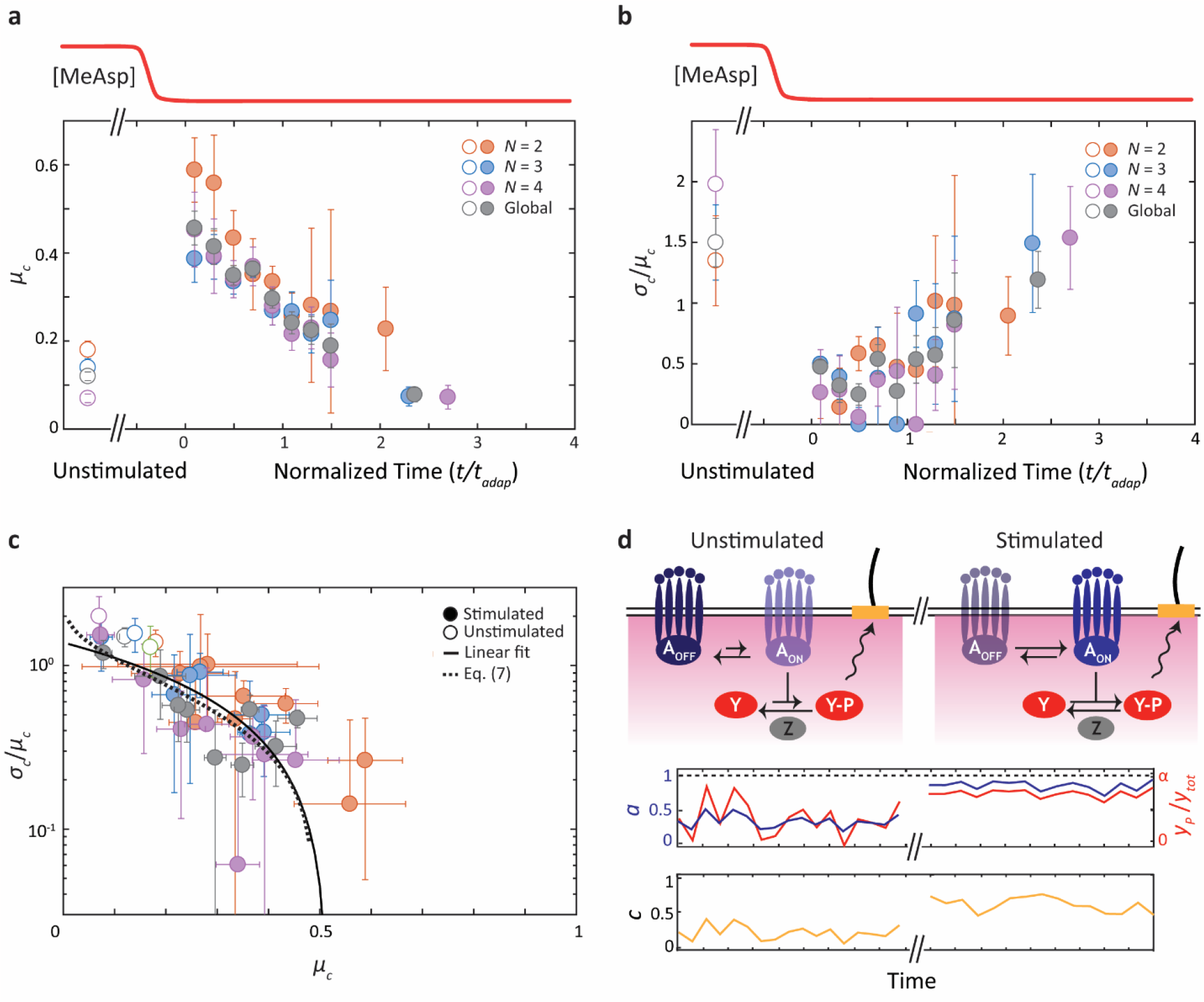
Reduction in motor bias fluctuations post stimulus due to kinase phosphorylation dynamics. **a**. Mean motor bias, *μ*_*c*_, vs. normalized time *t/t*_*adap*_ after step down stimulus (top schematic, red) determined separately for each number of flagella *N* (colored circles) and averaged across *N* (gray circles). **b**. Coefficient of variation in motor bias, *σ*_*c*_/*μ*_*c*_, after step down. Same color code as a. Data are pooled by cells with the same total number of flagella *N* = 2, 3, 4 (number of cells = 8, 9, 9). The pre-stimulus mean and coefficient of variation in motor bias were obtained from the analysis of the unstimulated cells in Fig. 2. **c**. Coefficient of variation in motor bias, *σ*_*c*_/*μ*_*c*_, plotted against the mean motor bias, *μ*_*c*_, for data from unstimulated and stimulated cells from a and b. Fits to a line (solid line) and to a model of network noise (dotted line; see Eq. (7)) are also shown. **d**. Simple model of network fluctuations in *E. coli*. (Left) In unstimulated cells at steady state, fluctuations in CheA activity (dark blue) produce bursts of CheY-P production (red) and high motor bias fluctuations (mustard). (Right) In stimulated cells, CheA activity is increases and shows reduced fluctuations as it approaches the kinetic ceiling. This also causes a reduction in [CheY-P] fluctuations and in motor bias fluctuations at high mean [CheY-P].

These results point to two interesting features. First, the mean CW bias shows an almost 300% increase compared to the pre-stimulus steady state immediately after the step down (**Figure 5a**). While the increase in CW bias is expected for a step-down stimulus, it is smaller in magnitude than predicted by existing models^54,55^. Given the strength of the stimulus (from 1 to 0 mM [MeAsp]), these models predict a CW bias close to 1; instead, the CW bias remains near 0.5. Second, fluctuations are significantly reduced immediately after the stimulus is applied, decreasing from *σ*_*c*_/*μ*_*c*_ = 1.5 pre-stimulus to as low as *σ*_*c*_/*μ*_*c*_ = 0.25 post-stimulus (**Figure 5b**). This value is lower than that measured in individual CheY* cells (**Figure 3b**), suggesting that fluctuations in the network are largely suppressed when driven to high CW bias. Noise suppression has previously been mentioned in the opposite limit^56^ when the network is driven to low activity by a step-up stimulus.

To summarize all of the experimental data, we plotted in **Figure 5c** the coefficient of variation vs. the mean CW bias for cells at steady state (open circles) and for stimulated cells (filled circles). All of the data fall on one continuous curve, which shows how network fluctuations vary across the accessible range of motor CW bias. Across all data sets, fluctuations uniformly exhibit the same behavior: a decreasing trend from high variance (*σ*_*c*_/*μ*_*c*_ > 1) at low CW bias, approaching zero at a CW bias of 0.5. For many biological networks, the relationship between the CV (or variance) and mean has proven useful in understanding the noise characteristics of the network^9,57^. For instance, Poisson-distributed noise is manifested by a CV decreasing as 1/mean^0.5^. Here, the trend can be fit to a line of form *C* (1 − *μ* / *μ*_*c*,max_), with *C*_0_ = 1.4 and *μ*_*c*,max_ ≈ 0.5, (solid line, **Figure 5c**). In the following discussion, we propose a model to interpret this trend and to understand the network noise characteristics they represent.

## Discussion

In this work we demonstrate a new approach for inferring fluctuations in the chemotaxis network based on the statistics of flagellar switching in multi-flagellated *E. coli* cells. One advantage of this approach is that its accuracy in quantifying network noise does not depend on temporal averaging, which limited previous measurements to long time scales (>10 s)^31,39–41^. Many recent measurements of network noise have been made through FRET imaging systems that require data collection and averaging over long periods of time, and thus sacrifice temporal resolution. Single flagellar tracking using high speed cameras has emerged as an alternative to measure network noise at short timescales^36^, and only very recently has revealed short time fluctuations (<1s)^58^. Our approach to estimating network noise relies on fluorescence-based observations of multiple flagellar motors making independent simultaneous “measurements” of the intracellular CheY-P level, and tuning their CW bias in response. This feature makes it possible to measure fluctuations in CW bias (and infer the underlying network noise) over time scales of individual runs and tumbles and to track their time evolution. As demonstrated in **Figure 4**, we can accurately determine the CV in CW bias over a ∼3-s time window (see **Materials and Methods**; this time resolution can in principle be improved further by pooling over a larger number of cells). In turn, studying cells as they undergo adaptation enables us to quantify network fluctuations across the accessible range of motor CW biases, as summarized in **Figure 5**. In **Figure 5c** the relationship between the CV and mean in CW bias provides a signature for the underlying *E. coli* chemotaxis signaling network noise, against which we can test various models. We thus sought a quantitative model to reproduce the noise characteristics we observed. The CW bias of every flagellar motor depends on the CheY-P concentration in an ultrasensitive switch-like manner (**Figure 1b, ii**)^25^, customarily modeled by a sigmoidal Hill equation of the form

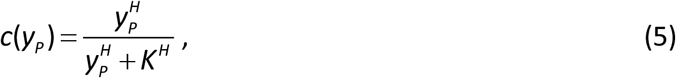

where *y*_*P*_ ≡ [CheY-P], *K* is the CheY-P concentration at 0.5 CW bias, and *H* is the Hill coefficient. (For cells at steady state, *K* ≈ 3.1 μM and *H* is expected to be ∼20^59^.) Due to this switch-like dependence, *c* is sensitive to small temporal fluctuations in [CheY-P]. CheY undergoes fast phosphorylation-dephosphorylation reactions (**Figure 1a**), which lead to number fluctuations in CheY-P (**Figure 1b, i**):

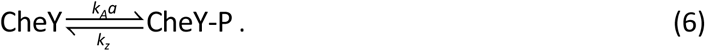

Here, the rate of CheY phosphorylation by the CheA-receptor complex is written as *k*_*A*_*a*, where *a* is the probability that CheA is in its active state and *k*_*A*_ is the maximum phosphorylation rate^60,61^, and *k*_*Z*_ is the rate of CheY-P dephosphorylation by the phosphatase CheZ. The intracellular [CheY-P] fluctuates due in part to the stochastic nature of the reactions in Eq. (6) and in part to fluctuations in the CheA kinase activity *a* itself^31,41^.

In **SI discussion**, we examine how different sources of noise can reproduce the observed behavior in CW bias fluctuations. Simple models incorporating CheY-P number fluctuations through Eq. (6) do not recapitulate the experimental trends. The main issue is that these models predict that CW bias fluctuations should to tend to zero as *c* approaches 1. This behavior occurs because *c*(*y*_*P*_), in Eq. (5), exhibits a plateau when [CheY-P] >> *K*, making it insensitive to fluctuations in [CheY-P] when CW bias is close to unity. However, as our data in **Figure 5c** shows, *σ*_*c*_/*μ*_*c*_ decreases to zero when CW bias = 0.5. Lele et al.^62^ previously carried out experiments ruling out the possibility of a plateau in *c* vs [CheY-P] at intermediate CW bias levels. This finding suggests that CheY-P fluctuations themselves must be suppressed when *c* = 0.5.

One possibility to account for this observation is that there exists a cap on the amount of CheY available for phosphorylation. For example, if such a cap limited the maximum CheY-P concentration to [CheY-P] ≈ *K*, then network fluctuations would be suppressed near a CW bias of 0.5 according to Eq. (5). To test this mechanism, we constructed a mutant *ΔcheZ* strain (“RB03”; see **Materials and Methods**) lacking the phosphatase CheZ, in which all the available CheY is expected to be phosphorylated. Replicating our flagellar imaging analysis for this strain resulted in a mean CW bias of 0.99 ± 0.01 (data not shown), higher than that observed following a step-down stimulus, ruling out such a mechanism. (The CV for this strain was 0.02; data not shown.) This result is consistent with estimates of the total concentration of CheY of ∼10 μM >> *K*^55^. An alternative possibility is that there exists a kinetic ceiling on the CheA-receptor complex, which limits the amount of CheY that can be phosphorylated at any one time. Since CheY-P undergoes dephosphorylation in the reaction scheme Eq. (6), the fraction of CheY that is phosphorylated must be less than one even when CheA is maximally active. Such a ceiling in the maximum [CheY-P] would lead to a reduction in [CheY-P] fluctuations.

To explore the latter mechanism quantitatively, we considered a comprehensive model taking into account fluctuations in CheY-P number and in the activity of the CheA-receptor complex during phosphorylation-dephosphorylation kinetics. To model CheA fluctuations, we allowed CheA to interconvert between active and inactive states, leading to a fluctuating activity *a*(*t*). Adapting the approach of Paulsson et al.^57,63^ used to model gene expression noise, we derived an analytical solution for the coefficient of variation in [CheY-P] in the presence of CheA fluctuations (see **SI Discussion**). Writing this expression in terms of CW bias gives

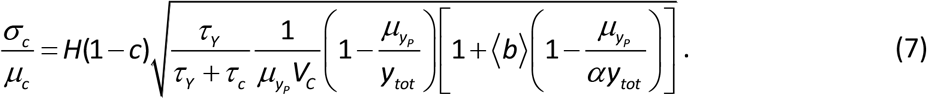

The appearance of the Hill coefficient *H* in Eq. (7) follows from the coupling of CheY-P fluctuations to those in CW bias *c* in Eq. (5). Here, *τ*_*c*_ is the characteristic timescale for motor switching and *τ*_*Y*_ =(*k*_*A*_*a* + *k*_*Z*_)^−1^ is that for CheY phosphorylation-dephosphorylation; the factor that depends on *τ*_*c*_ and *τ*_*y*_ accounts for temporal averaging that results from the different fluctuation timescales for motor switching and CheY reactions. The other factors inside the square root represent the number fluctuations in CheY-P; *y*_*tot*_ ≈ 10 μM is the total CheY concentration inside the cell, and *V*_*C*_ is the *E. coli* cell volume, assumed to be 1.4 fL^64^ 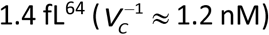. The factor in brackets specifically represents the contribution from CheA activity fluctuations. Here ⟨*b*⟩ denotes the number of new CheY-P generated during the time periods when CheA is active. The factor *α* ≡ *k*_*A*_ / (*k*_*A*_ + *k*_*Z*_) is the fraction of CheY phosphorylated when all CheA are active, i.e. when the activity *a* = 1.

This model recapitulates the features of the data well. **Figure 5c** shows a fit to Eq. (7) (dotted line) with parameters ⟨*b*⟩ = (6 ± 1) × 10^2^ and *α* = 0.32 ± 0.02 (*R*^2^ = 0.65; errors represent 95% confidence interval). Analogous to “burst size” in stochastic gene expression^65,66^, ⟨*b*⟩ increases number fluctuations in CheY-P, which are intrinsically low due to the high CheY copy number (see **SI Discussion**). The parameter *α* represents the fact that the contribution of CheA fluctuations to CheY-P noise must go to zero (as in **Figure 5c**) when *a* = 1, which corresponds to a mean CW bias of ∼0.5. (We note that the noise suppression previously mentioned by Colin et al.^56^ occurs in the opposite limit of *a* = 0.) The value of *α* points to a kinetic ceiling on CheA-receptor complexes of *k*_*A*_ = 0.47*k*_*Z*_, or ∼14 s^-1^ assuming a constant CheY-P dephosphorylation rate of *k*_*Z*_ = 30 s^-1^, within the reported range^54,60,61^. A maximum rate *k*_*A*_ = 14 s^-1^ is consistent with reported values of the CheA auto-phosphorylation rate, the rate-limiting step for CheY phosphorylation^33,54,60,61^. (We note that in the *ΔcheZ* strain, the kinetic ceiling on *k*_*A*_ would not cap [CheY-P] since *k*_*Z*_ = 0.)

Our observations suggest that network fluctuations are critical drivers for CW flagellar rotation/tumbling in *E. coli*. Previous studies^36,42,67^ proposed that waves of CheA activity could cause transient increases in [CheY-P]. Our data and model appear to be consistent with this mechanism, with the “burst size” ⟨*b*⟩ in Eq. (7) representing such waves in activity (see **Fig. 5d** for a schematic depiction). Our fit parameters suggest that several hundred CheY molecules are phosphorylated during such events, a not-insignificant fraction of the total number, estimated to be ∼8000^55^. We envision that transient increases in [CheY-P], driven by waves of phosphorylation by the receptor kinase complex^36^, increases the CW bias to generate tumbles. However, the amount of cellular [CheY-P] is subject to the constraints of CheA phosphorylation kinetics. When the cell experiences a network activating stimulus, and the receptor kinase complex is pushed to its maximum activity, the cellular [CheY-P] reaches its upper limit. This is manifested as high CW bias and suppressed fluctuations in CW bias.

Previous single-cell measurements of flagellar rotation by bead tracking^31,39^ and of CheA activity using fluorescent reporters^40,41^ have reported long-term noise in the chemotaxis network. This has largely been attributed to CheR-CheB methylation-demethylation dynamics^31,68^ and, recently, to new sources such as receptor clustering and other dynamics^40,41,67^. A direct comparison between these results and ours is difficult because of the differences in timescales between the measurements (3-40 s vs. 10-1000 s). Notably, burst-like noise similar to what we observe^36,42^ has been attributed to cooperative receptor state switching^67^. While our approach aligns with other recent work that extends noise measurements to fast timescales of the order of run and tumble durations, photobleaching of the dyes limits the duration of our traces and limits the overlap between the measurement timescales. For this reason, we have deliberately made our model for CheA activity fluctuations in Eq. (7) agnostic to the source of noise to allow for various possibilities. (We nevertheless note that the general form for CheA activity noise is similar to that used by Colin et al.^56^; see **SI Discussion** for more details.)

The noise characteristics of the chemotaxis network revealed in our measurements may have several functional consequences. First, network fluctuations may act to synchronize flagellar switching^36,37,42,67^, which was shown to make swimming behavior robust to variation in flagella number^37^. *E. coli* has also been proposed to exploit long-timescale network fluctuations to explore larger volumes of space^31,32^ and to increase drift speeds, enhancing migration along increasing attractant gradients^33–35^. In addition, our observation of a mean CW bias well below the midpoint of the *c* vs CheY-P curve at steady state is consistent with theory and simulations^45^ predicting that cells’ drift velocity along chemical gradients is maximized at low CW bias^47^, an idea supported by recent experiments^46^. Lastly, the kinetic ceiling in receptor kinase activity which restricts network fluctuations when the mean [CheY-P] increases (as depicted in **Fig. 5d**) may prevent CheY-P concentrations from saturating and leading to continuous and unproductive CW flagellar rotation/tumbling. Such behavior could result in increased *E. coli* swimming efficiency down steep decreases in attractant concentration, as likely found in natural environments. These findings point to network fluctuations tuned to maximize chemotactic drift.

## Supporting information

Supplementary Information (revised)

## Acknowledgements

We are thankful to Tom Kuhlman for guiding us through the process of molecular cloning, and to Howard Berg for providing bacterial strains. We thank members of the Chemla laboratory for scientific discussions and Christopher Rao for providing comments on the manuscript. Work in the Chemla and Golding labs is supported by National Science Foundation Physics Frontiers Center (PFC) “Center for the Physics of Living Cells” (CPLC) PHY-1430124. Work in the Golding lab is supported by the National Institutes of Health grant no. R35 GM140709 and the Alfred P. Sloan Foundation.

## Data availability

The datasets generated during and analyzed during the current study are available as an excel file included with this manuscript.

## Materials and Methods

### Microbiology

#### Cell preparation

Experiments were performed on the HCB1660 *E. coli* strain, a gift from the Berg lab^69^. This strain is considered wild-type for chemotaxis but is *ΔfliC* expressing FliC^S219C^ under the control of the P_araBAD_ promoter. The mutant FliC^S219C^ protein was specifically labelled with a cysteine reactive fluorescent dye, Alexa Fluor 532 C_5_ Maleimide (A10255, ThermoFisher Scientific) using the method first described by Turner et al.^69^ and later used by Mears et al.^37^. The strain referred to as CheY* is PM87, *ΔfliC ΔcheBYZ* expressing FliC^S219C^ and constitutively active CheY^D13K^, constructed by Mears et al.^37^. The strain referred to as RB03 (constructed for this work) is *ΔfliC ΔcheZ* expressing FliC^S219C^.

As described by Mears et al.^37^, for each experiment, a single colony from an LB, lysogeny broth, agar plate was inoculated into 1 mL tryptone broth (TB: 0.8% [wt/vol] NaCl, 1% Bacto Tryptone), and grown to saturation overnight (14-18 hours), shaking at 265 RPM at 30°C with the appropriate antibiotics. The overnight culture was diluted 100-fold into 12 mL TB, and grown to OD_600_ ∼0.4 to 0.5, with 0.01% [wt/vol] or 666 μM L-arabinose, shaking at 265 RPM at 30°C for ∼4.5 hours. The over-day culture was washed twice by slow centrifugation (1300 × *g*) and gently resuspended in 1 mL motility buffer (MB: 10 mM Phosphate Buffer [pH 7.0], 70 mM NaCl and 0.1 mM EDTA)^21^.

To label flagella fluorescently, we followed the protocol reported by Mears et al.^37^. An appropriate volume of the Alexa Fluor 532 C_5_ Maleimide dye was added to resuspended cells in 0.5 mL MB to a concentration of ∼0.01 mg/mL and incubated with slow rotation (∼10 RPM) at room temperature in the dark for 90 min. The labeled cells were gently resuspended in 1 mL MB. For trapping, cells were diluted 20-fold in trap motility buffer (TMB: 70 mM NaCl, 100 mM Tris-Cl, 2% [wt/vol] glucose, with 0.1 mM methionine and an oxygen-scavenging system [290 μg ml^−1^ pyranose oxidase and 65 μg ml^−1^ catalase]^38^) and injected into the flow chamber for the trapping experiment. When handling resuspended cells, at all times, we avoided pipetting, or we used wide orifice pipette tips to avoid shearing flagella.

1 mg of the dry Alexa Fluor 532 C_5_ Maleimide dye was dissolved in 250 μL water by vortexing and stored in aliquots at -20°C. The stock concentration for each aliquot was determined using UV-Visible Absorption Spectroscopy. Since this dye does not dissolve readily in water, we found it important to determine the concentration for each aliquot separately for optimal labeling.

### Construction of strains

The bacterial strains and plasmids used for this work are listed in **Table 1**. Oligonucleotides for generating mutations and creating plasmids are listed in **Table 2**. All primers were purchased from Integrated DNA Technologies (Coralville, Iowa). The *CheZ* chromosomal deletion was carried out using the standard λ Red recombinase system developed by Datsenko and Wanner^70^. The strain referred to as RB03 was created from HCB1613 (AW405, chemotaxis wild-type with a functional FliC deletion). To create the strain RB01, *CheZ* was replaced by a chloramphenicol resistance cassette with flanking FRT (Flp recombinase-recombination target) sites using the primers CheZ-pKDF and CheZ-pKDR with pKD3 as a template. This replacement was accomplished via the λ Red recombinase enzymes expressed from pKD46. The chloramphenicol cassette was eliminated via Flp recombinase expressed from pCP20 to obtain strain RB02. Finally, the strain RB03 was created by transforming strain RB02 with the plasmid pBAD33 that expresses FliC^S219C^. Standard molecular cloning techniques were used to purify plasmids and perform PCRs.

## Data Acquisition

### Optical trapping of individual *E. coli* with simultaneous fluorescence imaging

Experiments were performed with a dual-trap optical tweezers instrument combined with epi-fluorescence slim-field imaging, developed and described in detail in previous work by Mears et al.^37^. Briefly, the optical traps were created from a single 5-W, 1064-nm laser (YLR-5-1064-LP, IPG Photonics). Two trapping beams were generated by timesharing, intermittently deflecting the input beam between two positions with an Acousto-Optic Deflector, AOD (DTSXY-250-1064, AA Opto-Electronic). The two resulting beams were then tightly focused using a 60X, water immersion (1.2 NA) objective (Nikon, Tokyo, Japan) to create two optical traps. An identical objective was used to collect the transmitted trap light for position detection and for bright-field imaging of the cells using Koehler illumination from a blue LED. A bacterial cell could be captured and aligned between the two optical traps without impeding its rotatory motion. The light scattered by the trapped cell was collected onto a position sensitive photodetector and used to monitor cell motion, which was analyzed to determine runs and tumbles as described below and previously by Min et al.^38^.

Fluorescently labeled flagella of trapped cells were imaged using a 532-nm laser (TECGL-30, World Star Tech) aligned for slim-field illumination^71^, with the excitation beam diameter at the sample plane ∼10-15 μm. We strobed the fluorescence excitation and trap laser illumination out of phase with each other to minimize photobleaching^37^. The fluorescence illumination was strobed by using an Acousto-Optic Modulator, AOM (802AF1, IntraAction) to deflect the excitation light intermittently into the sample plane for a duration of 16-20 μs each cycle while the traps were off. An EMCCD camera (iXon3 860 EMCCD, Andor) recorded images in synchrony with the excitation pulse at the rate of 100 frames per second to capture focused images of the flagella as they rotated at a rate of ∼100 Hz. In this manner, movies of flagellar motion were recorded, and flagella waveforms were analyzed until the flagella photobleached, typically for 10-20 for unstimulated cells and 20-40 s for stimulated cells. Movies were saved using the Solis software (Andor) and processed using custom written code in MATLAB. Each movie was analyzed manually to count flagella and their transitions between CW and CCW rotating states.

### Motility assays combined with fluorescence imaging in trapped cells

Custom-built microfluidic chambers shown in **Figure 1c** and **4a** (preparation described in detail in Mears et al.^37^ and Min et al.^43^) were used to perform motility assays coupled with fluorescence imaging of labeled flagella on individual trapped cells. To prepare chambers, glass coverslips were sonicated in acetone, then rinsed in deionized water. This step was followed by rinsing in methanol and drying under nitrogen flow. The flow channels in the chamber were cut out from Nescofilm (Alfresa Pharma Corporation, Osaka, Japan) and bonded to glass coverslips as described previously^37,43^.

For measurements of free-swimming, or unstimulated, cells, the chamber contained two channels (**Figure 1c**). The upper channel contained trap motility buffer (TMB) only, while the lower channel contained *E. coli* cells in TMB. Both channels were continuously injected with the appropriate solutions using a syringe pump (PHD2000, Harvard Apparatus) with a linear flow rate of ∼30 μm/s. Cells were trapped in the lower channel and moved to the upper channel by displacing the chamber with respect to the traps using a motorized three-axis translational stage (ESP301; Newport, Irvine, California). Once the trapped cell was aligned between the two traps in the upper channel, the bright-field illumination was turned off and the 532-nm excitation illumination was turned on. Cell motion was observed by simultaneously recording the optical trap signal sensed by position sensitive photodetectors and the fluorescence signal from labelled flagella sensed by the EMCCD camera.

For measurements of stimulated cells undergoing chemotactic adaptation, the chamber contained three channels (**Figure 4a**). For a step-down stimulus, the top (“blank”) channel contained TMB only, the middle (“blank + attractant”) channel contained TMB, 1 mM *α*-methyl-DL-aspartate (MeAsp) and Rhodamine B (100 nM), and the bottom (“cell + attractant”) channel contained TMB, 1 mM MeAsp and *E. coli* cells. Fluorescence imaging of the Rhodamine B diffusion profile was used to characterize the gradient formed across the top and the middle channel, as described by Min et al.^43^. All three channels of the flow chamber were injected with their solutions at a linear flow rate of ∼50 μm/s. In a typical experiment, a swimming cell was first captured from the “cells + attractant” channel. The trapped cell was moved to the “blank + attractant” channel, aligned between the two traps, and oriented along the direction of the flow. Pre-stimulus optical trap signal of the swimming cell was recorded for ∼200 s (without any fluorescence imaging of flagella). The trapped cell was then moved at the speed 100 μm/s to the “blank” channel so that it experienced a step down in MeAsp concentration. While the entire move took 10 s, the interface between channels where the step down occurred was crossed in ∼4 s (where the interface is defined as the distance from 10% to 90% of the chemoattractant concentration, as shown in Figure S1 in Min et al.^43^). *t* = 0 was defined as the end of this move. During the move, the bright field illumination was turned off and the 532-nm excitation illumination was turned on. The chemotactic response of the captured cell was recorded in the “blank” channel by simultaneously recording the optical trap signal and the fluorescence signal from labelled flagella. While the trap signal post-stimulus was recorded for ∼400 s, the EMCCD signal was recorded only for ∼30-40 s until the flagella photobleached.

## Raw Data Analysis

### Run-tumble analysis for optical trap data

Cell motion was detected from imaging the optical trap light scattered by the cell body onto position-sensitive photodetectors, as described previously by Min et al.^38^. This optical trap signal was analyzed for runs and tumbles. Briefly, runs corresponded to oscillatory signals due to cell body roll at a frequency ∼10 Hz, while tumbles corresponded to erratic trap signals with a frequency <1 Hz. We used custom-written MATLAB code incorporating a wavelet analysis to identify the peak body roll frequency at every time point of each data trace and to assign runs and tumbles by specifying an appropriate frequency threshold^38^. An example trace of the trap signal and the corresponding run-tumble assignment is shown in **Figure 1c** and **d** in earlier work by Mears et al.^37^.

### Image analysis for fluorescence data

Images acquired using the high-speed fluorescence imaging described above were adjusted for contrast to best visualize the flagella. Each movie was played in slow motion and Clockwise (CW)/Counter Clockwise (CCW) states were assigned manually to each flagellum over a time window of 100 ms (i.e. 10 frames). As described previously^21,37^, the CCW/CW rotation state of the flagella can be visually identified from their different helical waveforms, commonly referred to as ‘normal’, ‘semi-coiled’, ‘curly-1’ and ‘curly-2’. CCW-rotating flagella exclusively adopt the ‘normal’ waveform and CW-rotating flagella adopt the ‘semi-coiled’ and ‘curly-1’ waveforms the majority of the time. Waveform transitions take less than ∼0.5 s to propagate through the flagella.

## Modeling and Analysis

### Extracting CW bias fluctuations from flagellar rotation data

In flagellar imaging movies of individual cells, we tracked the CW/CCW state of each flagellum— hence the number of CW flagella in time—and determined *p*_*N*_ (*n*_*cw*_), the probability that *n*_*cw*_ out of a total of *N* flagella rotate CW, where *n*_*cw*_ = 0, 1… *N*. From first principles^72^, we can relate *p*_*N*_ (*n*_*cw*_) to the instantaneous CW bias of the motor, *c*, the probability that a motor rotates CW, through Eq. (1), where *p*(*c*) is the distribution in motor bias *c. p*_*N*_ (*n*_*cw*_ |*c*), the conditional probability that *n*_*cw*_ out of *N* flagella are CW given a certain bias *c*, is given by the binomial distribution in Eq. (2), since as shown in previous work by Mears et al.^37^, the flagellar motors switch independently.

We can thus use Eq. (1) and (2) to extract information on the motor bias distribution *p*(*c*) from measurements of *p*_*N*_ (*n*_*cw*_). In general, this integral equation cannot be solved by a unique solution *p*(*c*). Instead, a finite number of moments of *p*(*c*) can be determined, which increases as the number of flagella *N* increases. Using Eq. (1) and (2), one can relate moments of *p*_*N*_ (*n*_*cw*_) to those of *p*(*c*). For the first two moments, it can be shown that:

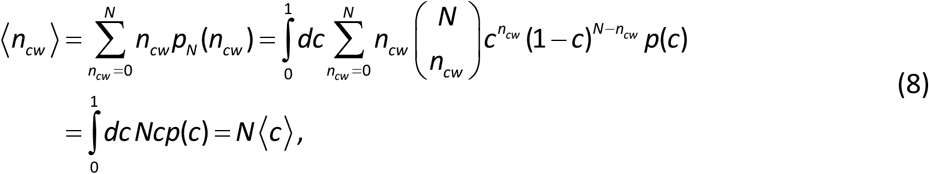

and

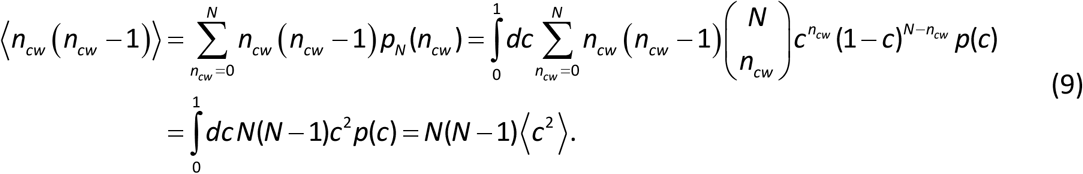

Eqs. (8) and (9) can be manipulated to give Eqs. (3) and (4) for the mean and variance of the CW bias in the text. We note that Eq. (9) shows that ⟨*c*^2^⟩ can be determined only from cells with *N* ≥ 2 flagella. The first two moments are generally adequate for our purposes to assess fluctuations in the network, and we analyzed cells with *N* ≥ 2. In principle, this approach can be extended to determine higher moments (of order *n* ≥ 3), which reveal finer features of the CW bias distribution *p*(*c*) (e.g. skew, kurtosis, etc.). We chose not to determine these higher moments due to the limited amount of cells with *N* ≥ *n* flagella.

Since the form of the distribution *p*_*N*_ (*n*_*cw*_) depends on the number of flagella, we analyzed cells with different numbers of flagella *N* separately, as shown in **Figure 2a** and **d**. For each population of cells, we determined the mean and standard deviation in CW bias and estimated their errors by bootstrapping, where individual cell time traces were resampled with replacement to generate different *p*_*N*_ (*n*_*cw*_) for each bootstrap trial. We also determined global values for these parameters across the population of cells with different *N* by carrying out an inverse-variance weighted average over their values for each *N*. The errors were determined using the customary expression for the standard error of a weighted mean.

Finally, we also used distributions *p*_*N*_ (*n*_*cw*_) obtained from each individual cell to determine the mean and standard deviation in CW bias from each cell (**Figure 3**). The errors in these parameters were estimated using bootstrapping. Here, 0.5-s long time windows of individual cell time traces of *n*_*cw*_ were resampled with replacement to construct traces and corresponding *p*_*N*_ (*n*_*cw*_) for each bootstrapping trial. A 0.5-s window duration was chosen based on the average autocorrelation time for individual cell *n*_*cw*_ time traces, determined to be ∼0.3 s across *N* (data not shown). Time windows of duration of 0.5 s, about twice as long as the autocorrelation time, are sufficiently long to be uncorrelated and suitable for the purposes of bootstrapping.

### Variance decomposition

The variance in CW bias measured over a population of cells contains contributions from the single-cell CW bias variance and from cell-cell differences in mean CW bias. We decomposed the variance over the population (the set {cell}) to determine the intra- and inter-cell contributions, using the Law of Total Variance^73^:

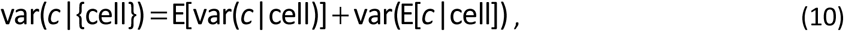

where E[…] is the expectation value and var is the variance. Based on established interpretations of Eq. (10), the first term on the right-hand side represents the contribution of the average single-cell variance in CW bias, which is determined from the mean of the individual variances in CW bias for each cell for a given *N*. The second term on the right hand side is the contribution of the cell-to-cell variation in CW bias, which is determined from the variance in the mean CW bias for each cell for a given a *N*.

### Estimating CW bias fluctuations for stimulated cells

The flagellar imaging data for trapped cells experiencing a step-down in MeAsp were also analyzed to record the number of CW flagella with time. Since the motor bias varies with time during adaptation, each individual cell time trace was divided into non-overlapping windows the duration of which was determined from the cell’s adaptation time. For each cell, the collected optical trap signal was analyzed for runs and tumbles, and the tumble bias was calculated from the fraction of time the cell tumbled over a 10-s sliding time window. The time trace of the tumble bias was fit to the following expression:

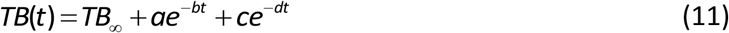

where *a, b, c* and *d* are fitting parameters, and *TB*_*∞*_ is the tumble bias of the cell when it has adapted to the stimulus. The adaptation time was approximated as *t*_*adap*_ =ln(2)/ *b* following a procedure used by Min et al.^43^ An example fit to Eq. (11) is shown in **Supplementary Figure S1**.

Windows of duration 0.2*t*_*adap*_ were selected starting at the time when the trapped cell was moved to the top channel, *t* = 0, until *t* = 1.6*t*_*adap*_, where *t*_*adap*_ for our dataset was found to be ∼13 s. A final window of variable length between 0.5*t*_*adap*_ and 2.5*t*_*adap*_ was created based on the number of data points left in the trace. Data for number of CW flagella *n*_*cw*_ from multiple cells with the same total number of flagella *N* were pooled in each time window. The time-dependent histogram *p*_*N*_ (*n*_*cw*_, *t*) was constructed and the time-dependent mean and standard deviation in CW bias was determined for each time window. The 0.2*t*_*adap*_ time windows, corresponding to a 2.6-s duration on average, were sufficiently short so that the average tumble bias was constant to within ∼10-20%, but long enough to enable a reliable estimate of the mean and CV in CW bias given the data set size. The error was calculated by bootstrap resampling as before, over each time window. We calculated the mean and standard deviation in CW bias globally across all *N* for each time window as described previously.

