## Supplementary Information (revised) for "Flagellar dynamics reveal fluctuations and kinetic limit in the *Escherichia coli* chemotaxis network"

**Contents:**

Supplementary Figure S1

Supplementary Discussion

### Supplementary Figures

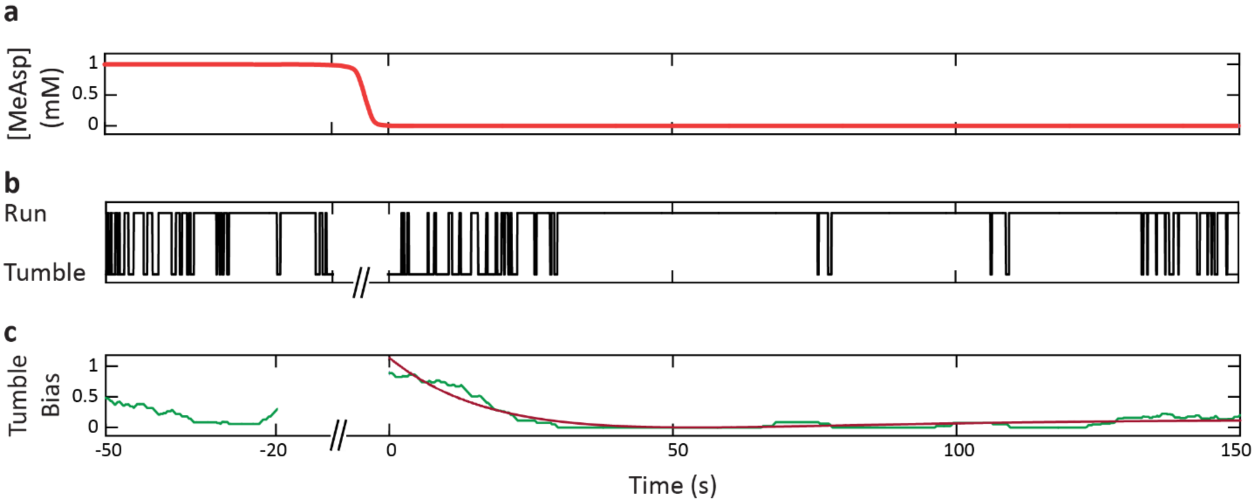


**Supplementary Figure S1. Run-tumble response of a cell to a step-down stimulus. a.** Schematic of MeAsp concentration over time, showing the step down from 1 to 0 mM where $t = 0$ is the time when the cell finished moving to the channel with no MeAsp. **b.** Run-tumble binary time trace of a representative cell. Data are not collected during the move from the middle to the top channel. **c.** Tumble bias of the cell (green), determined at each time point from the fraction of time tumbling over a 10-s moving window, and fit to bi-exponential function (red) Eq. (11).

### Supplementary Discussion

#### Analytical model of fluctuations in number of CW flagella

In our assay, we determine the instantaneous number of flagellar motors that rotate CW vs CCW, and use fluctuations in these numbers to infer noise in the underlying chemotaxis network. We use a stochastic chemical kinetics approach to model these fluctuations analytically. Following previous work[1], we consider that each flagellar motor can interconvert between rotational states according to the following reaction, given a CW bias *c*:

$$\mathrm{Motor_{CCW}} \underset{k(1-c)}{\overset{kc}{\rightleftharpoons}} \mathrm{Motor_{CW}}\,.$$

Here, $1/kc$ and $1/k(1\text{-}c)$ are the CCW and CW time intervals, respectively, and $k$ is the characteristic switching rate constant. We then use a chemical master equation (CME) approach to determine fluctuations in the number of CW flagellar motors $n_{cw}$ for this reaction. The individual reactions in terms of the numbers in each rotational state are:

Motor switch from CCW to CW: $\quad n_{cw} \xrightarrow{kc(N-n_{cw})} n_{cw}+1$

Motor switch from CW to CCW: $\quad n_{cw} \xrightarrow{k(1-c)n_{cw}} n_{cw}-1$

where $N$ is total number of flagellar motors. The motor switching transitions are assumed to be independent of each other.

Following the approach of McQuarrie and Gillespie[2], the CME for the above can be solved analytically to yield the following steady-state mean and variance in the number of CW motors,

$$\langle n_{cw}\rangle = cN\,, \tag{S1}$$

$$\sigma_{n_{cw}}^2 = Nc(1-c) = \langle n_{cw}\rangle\left(1-\frac{\langle n_{cw}\rangle}{N}\right). \tag{S2}$$

Eqs. (S1) and (S2) can be combined to obtain the coefficient of variation,

$$\frac{\sigma_{n_{cw}}^2}{\langle n_{cw}\rangle^2} = \frac{1-c}{\langle n_{cw}\rangle}\,, \tag{S3}$$

which has the expected form for a binomial-distributed process. Eq. (S3) represents the fluctuations intrinsic to switching of $N$ independent flagellar motors, but does not include contributions from noise in CW bias, $c$, which is implicitly assumed to be constant. To account for CW bias fluctuations, additional terms must be added to Eq. (S3), which must collectively sum to the squared coefficient of variation in CW bias $\sigma_c^2 / \mu_c^2$. (This can be seen by comparing Eqs. (S1) and (S2) to Eqs. (3) and (4), respectively, in the main text).

**Analytical model of fluctuations in number of CW flagella with CheY noise**

To model CW bias fluctuations analytically, we considered sources of noise upstream in the signaling network that could affect downstream behavior. While various models[3,4] describe a wide range of reactions in the chemotactic signaling network, we chose the simplest possible scenario to verify that it could recapitulate the essential features of our experimental results.

The motor CW bias $c$ depends on the intracellular CheY-P concentration in an ultrasensitive switch-like manner, customarily modeled by a sigmoidal Hill equation[5],

$$c(y_P) = \frac{y_P^H}{y_P^H + K^H}, \tag{S4}$$

where $y_P \equiv$ [CheY-P], $K$ is the CheY-P concentration at 0.5 CW bias, and $H$ is the Hill coefficient. (For simplicity, we assume that the parameters $H$ and $K$ are constant over the timescales of our measurements. This assumption is justified since the measurement timescales are significantly shorter than the those of flagellar motor remodeling known to influence $K$ and $H$[6,7].) Due to this switch-like dependence, the CW bias is sensitive to temporal fluctuations in [CheY-P]. We first considered number fluctuations in CheY-P due to CheY phosphorylation and dephosphorylation reactions. Here,

$$\text{CheY} \underset{k_Z}{\overset{k_A a}{\rightleftharpoons}} \text{CheY-P},$$

where $k_A a$ is the rate of phosphorylation of CheY by the CheA-receptor complex and $k_Z$ is the rate of dephosphorylation of CheY-P by the phosphatase CheZ. As is standard practice[3,8], the activity $a$ of the CheA-receptor complex is defined as the probability that it is in the active state ($0 \le a \le 1$), and $k_A$ as the maximum phosphorylation rate.

Eq. (S4) couples the phosphorylation-dephosphorylation reactions to the motor CCW/CW switching transitions, which results in CheY-P noise contributing to fluctuations in CW bias. To determine this noise contribution, we use a CME approach to calculate number fluctuations for all molecular species. The individual reactions for each species are:

Motor switch from CCW to CW: $n_{cw} \xrightarrow{kc(N-n_{cw})} n_{cw}+1$

Motor switch from CW to CCW: $n_{cw} \xrightarrow{k(1-c)n_{cw}} n_{cw}-1$

Phosphorylation of CheY: $n_{Yp} \xrightarrow{k_A a(N_Y^{tot}-n_{Yp})} n_{Yp}+1$

Dephosphorylation of CheY-P: $n_{Yp} \xrightarrow{k_Z n_{Yp}} n_{Yp}-1$

where $n_{Yp}$ is number of CheY-P molecules, and $N_Y^{tot}$ is the total number of (unphosphorylated and phosphorylated) CheY molecules in the cell, assumed to be fixed.

Paulsson[9,10] previously developed a systematic formalism for calculating number fluctuations in gene expression, which we adapted for the reaction scheme above. The steady-state mean number of CheY-P is easily shown to be

$$\frac{\langle n_{Yp}\rangle}{N_Y^{tot}} = f_{Yp} = \frac{k_A a}{k_A a + k_Z}, \tag{S5}$$

where $f_{Yp}$ is the fraction of total CheY that is phosphorylated. The mean number of CW flagella is given by the same expression as Eq. (S1). For number fluctuations, Paulsson provides the following general solution[10]:

$$\frac{\sigma_{Yp}^2}{\langle n_{Yp}\rangle^2} = \frac{1}{\langle n_{Yp}\rangle H_{YY}},$$

and

$$\frac{\sigma_{n_{cw}}^2}{\langle n_{cw}\rangle^2} = \frac{1}{\langle n_{cw}\rangle H_{cc}} + \frac{\sigma_{Yp}^2}{\langle n_{Yp}\rangle^2}\left(\frac{H_{cY}}{H_{cc}}\right)^2 \frac{H_{cc}/T_c}{H_{cc}/T_c + H_{YY}/T_Y},$$

where the logarithmic gains $H_{ij}$ and times $T_i$ are defined in Eq. (S5) and (S6) in Ref. [9]. The first term in the expression for the CV in number of CW flagella represents the intrinsic number fluctuations in CW-rotating flagella from the switching reaction, while the second represents the noise contribution from CheY-P fluctuations. In the second term, the ratio of gains $H_{ij}$ determines how much CheY-P fluctuations couple to those in the number of CW flagella, and the factor containing the times $T_i$ accounts for the time averaging that results from the different fluctuation timescales.

Calculating the gains $H_{ij}$ and times $T_i$ according to Ref. [9], the coefficient of variation in number of CW flagella simplifies to

$$\frac{\sigma_{n_{cw}}^2}{\langle n_{cw}\rangle^2} = \frac{1-c}{\langle n_{cw}\rangle} + \frac{\sigma_{Yp}^2}{\langle n_{Yp}\rangle^2} H^2 (1-c)^2 \frac{\tau_Y}{\tau_Y + \tau_c}, \tag{S6}$$

where $\tau_c = k^{-1}$ is the characteristic timescale for motor switching and $\tau_Y = (k_A a + k_Z)^{-1}$ is that for CheY phosphorylation-dephosphorylation. The appearance of the Hill coefficient $H$ follows from the coupling of CheY-P fluctuations to those in CW bias $c$ in Eq. (S4). We note that the first term is identical to Eq. (S3); it follows that the second term corresponds to the coefficient of variation in CW bias. In other words,

$$\frac{\sigma_c}{\mu_c} = H(1-c)\sqrt{\frac{\tau_Y}{\tau_Y + \tau_c}} \frac{\sigma_{Yp}}{\langle n_{Yp}\rangle}. \tag{S7}$$

We can compare this model to the experimental data in **Figure 5c**. Based on the range of values for the rate constants $k_A$ and $k_Z$ in the literature[3,4], we expect the characteristic time scale $\tau_y$ to be ~0.025 s. Similarly, the motor switching time $\tau_c \approx 0.1$ s, as previously reported[1]. Thus, assuming that $H = 18$, a value measured by Yuan and Berg[11], a CV in [CheY-P] of ~0.23 reproduces the observed CV in CW bias for cells at steady-state ($\mu_c \approx 0.12$). This estimate of CheY-P fluctuations is consistent with a theoretically predicted lower bound of 20% of the mean[12]. Unfortunately, Eq. (S7) also predicts that CW bias fluctuations should tend to zero as $c$ approaches 1. This behavior occurs because $c(y_P)$, in Eq. (S4), exhibits a plateau when [CheY-P] >> $K$, making it insensitive to fluctuations in [CheY-P] when CW bias is close to unity. Our data in **Figure 5c** show that $\sigma_c/\mu_c$ decreases to zero when CW bias = 0.5. Lele et al.[13] previously carried out experiments ruling out the possibility of a plateau in $c$ vs [CheY-P] at intermediate CW bias levels. This disagreement between model and data suggests that CheY-P fluctuations themselves must be suppressed when $c = 0.5$.

We note that another prediction of the above model is that the coefficient of variation in the number CheY-P molecules has the form

$$\frac{\sigma_{Yp}^2}{\langle n_{Yp}\rangle^2} = \frac{1-f_{Yp}}{\langle n_{Yp}\rangle},$$

where $f_{Yp}$ is defined in Eq. (S5). This expression predicts a CV in [CheY-P] an order of magnitude smaller than that needed to match the experimental data, 0.23, as described above. Fundamentally, the phosphorylation-dephosphorylation reactions alone generate low number fluctuations CheY-P due to the large total number (~8000) of CheY molecules in the cell[14].

**Analytical model of fluctuations in number of CW flagella with CheA noise**

To model the key features of the experimental data in **Figure 5c**, we considered additional sources of noise that can contribute to the two-species model above. One such source is CheA activity in phosphorylation-dephosphorylation kinetics. Although $k_A$ and $k_Z$ are assumed to be fixed rate constants in the reaction scheme above, the activity $a$ is expected to fluctuate over time[15,16], which can couple noise in CheY-P number. To model fluctuations in $a$, we devised a simple model in which CheA interconverts between active (ON) and inactive states (OFF) with rates $\lambda_A^{\pm}$,

$$\mathrm{CheA_{OFF}} \underset{\lambda_A^-}{\overset{\lambda_A^+}{\rightleftharpoons}} \mathrm{CheA_{ON}}.$$

With this additional noise source, the individual reactions for the molecular species are given by:

| | |
|---|---|
| Activation of CheA: | $n_A \xrightarrow{\lambda_A^+ (N_A^{tot} - n_A)} n_A + 1$ |
| Deactivation of CheA: | $n_A \xrightarrow{\lambda_A^- n_A} n_A - 1$ |
| Phosphorylation of CheY: | $n_{Yp} \xrightarrow{\kappa_A n_A (N_Y^{tot} - n_{Yp})} n_{Yp} + 1$ |
| Dephosphorylation of CheY-P: | $n_{Yp} \xrightarrow{k_Z n_{Yp}} n_{Yp} - 1$ |
| Motor switch from CCW to CW: | $n_{cw} \xrightarrow{kc(N - n_{cw})} n_{cw} + 1$ |
| Motor switch from CW to CCW: | $n_{cw} \xrightarrow{k(1-c) n_{cw}} n_{cw} - 1$ |

where $n_A$ is the number of active CheA molecules, and $N_A^{tot}$ is the total number of CheA molecules in the cell, assumed to be fixed. Here, we made explicit the dependence of the phosphorylation rate on the number of active CheA, $n_A$, defining $\kappa_A$ such that $\kappa_A n_A = k_A a$, with the activity $a = n_A / N_A^{tot}$ and the maximum phosphorylation rate $k_A = \kappa_A N_A^{tot}$.

Number fluctuations in this set of coupled reactions can again be solved using the formalism developed by Paulsson[9,10]. The steady-state mean number of active CheA is

$$\frac{\langle n_A \rangle}{N_A^{tot}} = a = \frac{\lambda_A^+}{\lambda_A^+ + \lambda_A^-}, \tag{S8}$$

with the mean numbers of CW flagella and CheY-P given by Eqs. (S1) and (S5), as before. The number fluctuations for the different molecular species are:

$$\frac{\sigma_A^2}{\langle n_A \rangle^2} = \frac{1}{\langle n_A \rangle H_{AA}},$$

$$\frac{\sigma_{Yp}^2}{\langle n_{Yp} \rangle^2} = \frac{1}{\langle n_{Yp} \rangle H_{YY}} + \frac{\sigma_A^2}{\langle n_A \rangle^2} \left( \frac{H_{YA}}{H_{YY}} \right)^2 \frac{H_{YY}/T_Y}{H_{YY}/T_Y + H_{AA}/T_A},$$

and

$$\frac{\sigma_{n_{cw}}^2}{\langle n_{cw} \rangle^2} = \frac{1}{\langle n_{cw} \rangle H_{cc}} + \frac{\sigma_{Yp}^2}{\langle n_{Yp} \rangle^2} \left( \frac{H_{cY}}{H_{cc}} \right)^2 \frac{H_{cc}/T_c}{H_{cc}/T_c + H_{YY}/T_Y}$$
$$+ \frac{\sigma_A^2}{\langle n_A \rangle^2} \left( \frac{H_{cY}}{H_{cc}} \right)^2 \left( \frac{H_{YA}}{H_{YY}} \right)^2 \frac{H_{YY}/T_Y}{H_{cc}/T_c + H_{YY}/T_Y} \frac{H_{YY}/T_Y}{H_{YY}/T_Y + H_{AA}/T_A} \frac{H_{cc}/T_c}{H_{cc}/T_c + H_{AA}/T_A}.$$

Calculating the gains $H_{ij}$ and times $T_i$, these expressions simplify. The coefficient of variation in CheA becomes

$$\frac{\sigma_A^2}{\langle n_A \rangle^2} = \frac{1-a}{\langle n_A \rangle}, \tag{S9}$$

expected for a binomial-distributed process. A key feature of this type of model is that the CV in CheA decreases to zero as the kinase activity approaches 1. (We note that Colin et al.[16] quantitatively describe CheA fluctuations using a model of similar form as Eq. (S9).) The coefficient of variation in CheY-P simplifies to

$$\frac{\sigma_{Yp}^2}{\langle n_{Yp} \rangle^2} = \frac{1-f_{Yp}}{\langle n_{Yp} \rangle} + \frac{\sigma_A^2}{\langle n_A \rangle^2} (1-f_{Yp})^2 \frac{\tau_A}{\tau_A + \tau_Y}, \tag{S10}$$

where $\tau_A = (\lambda_A^+ + \lambda_A^-)^{-1}$ is the characteristic relaxation timescale for CheA activation-deactivation. Finally, the coefficient of variation in number of CW flagella becomes

$$\frac{\sigma_{n_{cw}}^2}{\langle n_{cw}\rangle^2}=\frac{1-c}{\langle n_{cw}\rangle}+\frac{\sigma_{Yp}^2}{\langle n_{Yp}\rangle^2}H^2(1-c)^2\frac{\tau_Y}{\tau_Y+\tau_c}+\frac{\sigma_A^2}{\langle n_A\rangle^2}H^2(1-c)^2(1-f_{Yp})^2\frac{\tau_c}{\tau_Y+\tau_c}\frac{\tau_A}{\tau_A+\tau_Y}\frac{\tau_A}{\tau_A+\tau_c}.$$

We note that in the absence of fluctuations in CheY and CheA, Eq. (S3) is recovered.

Substituting Eqs. (S9) and (S10) gives the following expression

$$\begin{aligned}\frac{\sigma_{n_{cw}}^2}{\langle n_{cw}\rangle^2}&=\frac{1-c}{\langle n_{cw}\rangle}+\frac{1-f_{Yp}}{\langle n_{Yp}\rangle}H^2(1-c)^2\frac{\tau_Y}{\tau_Y+\tau_c}\\&+\frac{1-a}{\langle n_A\rangle}H^2(1-c)^2(1-f_{Yp})^2\frac{\tau_Y}{\tau_Y+\tau_c}\frac{\tau_A}{\tau_A+\tau_Y}\frac{\tau_A+\tau_c+\tau_c\tau_A/\tau_Y}{\tau_A+\tau_c}.\end{aligned}\tag{S11}$$

The second and third terms in Eq. (S11) represent the contributions to CW bias fluctuations from noise in CheY-P number and CheA activity, respectively. Given Eq. (S11), we write the following expression for the coefficient of variation in CW bias:

$$\frac{\sigma_c}{\mu_c}=H(1-c)\sqrt{\frac{\tau_Y}{\tau_Y+\tau_c}\frac{1-f_{Yp}}{\langle n_{Yp}\rangle}\left[1+\frac{\langle n_{Yp}\rangle}{\langle n_A\rangle}(1-a)(1-f_{Yp})\frac{\tau_A}{\tau_A+\tau_Y}\frac{\tau_A+\tau_c+\tau_A\tau_c/\tau_Y}{\tau_A+\tau_c}\right]}.$$

From the steady-state solutions for the means of the molecular species, Eqs. (S5) and (S8), we obtain, after some algebra, a simpler expression

$$\frac{\sigma_c}{\mu_c}=H(1-c)\sqrt{\frac{\tau_Y}{\tau_Y+\tau_c}\frac{1-f_{Yp}}{\langle n_{Yp}\rangle}\left[1+\langle b\rangle\left(1-\frac{f_{Yp}}{\alpha}\right)\right]},\tag{S12}$$

where we define $\alpha\equiv k_A/(k_A+k_Z)$ and

$$\langle b\rangle\equiv\frac{N_Y^{tot}}{N_A^{tot}}k_A\tau_A\frac{\tau_Y}{\tau_A+\tau_Y}\frac{\tau_A+\tau_c+\tau_A\tau_c/\tau_Y}{\tau_A+\tau_c}.\tag{S13}$$

We note that the first factor outside the brackets in Eq. (S12) is the coefficient of variation in the absence of CheA fluctuations. The second, bracketed factor describes how CheA noise increases CheY-P number fluctuations. In analogy with transcriptional bursting[17,18], the parameter $\langle b\rangle$ in Eq. (S13) represents the "burst size", the number of new CheY-P molecules generated per CheA within its active lifetime. The other parameter, $\alpha$, describes the effect of a kinetic ceiling of CheA on CheY-P number fluctuations. $f_{Yp}/\alpha$ is the number of CheY-P molecules when all CheA-receptor complexes are active ($a$ = 1), i.e. when the system phosphorylates at its maximum possible rate. When CheA-receptor complexes attain this kinetic ceiling, the added

CheY-P fluctuations due to CheA noise drops to zero in Eq. (S12). **Figure 5c** shows a comparison between this model and the experimental data. In this figure, Eq. (S12) is plotted vs CW bias by first dividing the number of CheY-P molecules $\langle n_{Y_P} \rangle$ by the *E. coli* cell volume $V_C$ (assumed to be 1.4 fL[19]) to convert it into a concentration $y_P \equiv$ [CheY-P], and then using Eq. (S4) to convert the CheY-P concentration into CW bias *c*.
